# DEAD-box RNA Helicases Act as Nucleotide Exchange Factors for Casein Kinase 2

**DOI:** 10.1101/2021.12.20.473452

**Authors:** Edoardo Fatti, Alexander Hirth, Andrea Švorinić, Matthias Günther, Cristina-Maria Cruciat, Gunter Stier, Sergio P. Acebron, Dimitris Papageorgiou, Irmgard Sinning, Jeroen Krijgsveld, Thomas Höfer, Christof Niehrs

**Author notes:** Equal contribution (Edoardo Fatti, Alexander Hirth, Andrea Švorinić, Matthias Günther).

## Abstract

DDX RNA helicases promote RNA processing but DDX3X is also known to activate casein kinase 1 ε (CK1ε). Here we show that not only is protein kinase stimulation a latent property of other DDX proteins towards CK1ε, but that this extends to casein kinase 2 (CK2α2) as well. CK2α2 enzymatic activity is stimulated by a variety of DDX proteins and we identify DDX1/24/41/54 as physiological activators required for full kinase activity *in vitro* and in *Xenopus* embryos. Mutational analysis of DDX3X reveals that CK1 and CK2 kinase stimulation engages its RNA binding-but not catalytic motifs. Mathematical modelling of enzyme kinetics and stopped-flow spectroscopy converge that DDX proteins function as nucleotide exchange factor towards CK2α2 that reduce unproductive reaction intermediates and substrate inhibition. Our study reveals protein kinase stimulation by nucleotide exchange as a new principle in kinase regulation and an evolved function of DDX proteins.

## Main Text

DEAD box (DDX) proteins are a large family of RNA helicases that play key roles in a vast array of biological processes^1,2^. They unwind RNA structures and dissociate RNA-protein complexes in reactions fuelled by ATP hydrolysis but the function and biological role of most of the 44 human DDX proteins remains elusive. Among DDX helicases, DDX3X has an unconventional role during Wnt signalling, where it acts as specific activator of casein kinase 1ε (CK1ε; *CSNK1E*)^3,4^, consistent with its role in Wnt-dependent *Xenopus* embryonic axis formation^3^ and Wnt-driven cancers^5-7^. Casein kinase activation by DDX3X is evolutionary conserved since the *Neurospora* DDX3X homolog FRH activates CK1 in circadian clock regulation^8^.

It remained curious why an RNA helicase should stimulate a protein kinase and left unanswered what the underlying mechanism is. Notably, we observed that other, randomly selected DDX members also stimulated CK1ε activity *in vitro*^*3*^. Could kinase stimulation by DDX helicases be a more widespread phenomenon? If so, what is the underlying mechanism?

We here show that casein kinase 2 (CK2α2), which belongs to a different kinase family than CK1, is another DDX-regulated kinase for which we identified DDX1, -24, -41, and -54 as physiological activators, i.e. required for full kinase activity. Elucidating the mechanism, we carried out a detailed structure-function analysis with DDX3X and found that kinase stimulation resides in the helicase core domain and engages RNA binding-but not catalytic motifs. Extensive enzyme kinetics, mathematical modelling, and stopped-flow spectroscopy converge that DDX proteins function as nucleotide exchange factors (NEF) that reduce unproductive reaction intermediates and substrate inhibition. This notion is highly unexpected and has potentially wide-ranging implications.

### Several DDX proteins stimulate CK1ε *in vitro* but *in vivo* the requirement for DDX3X is specific

We first followed up on the observation, that randomly selected DDX members stimulate CK1ε activity *in vitro*^*3*^ by monitoring DDX binding and kinase stimulation. In pulldown experiments, DDX3X as well as six DDX proteins (DDX6, -20, -39A, -41, -50, -56) tested previously^3^, bound purified CK1α(*CSNK1A*), CK1δ (*CSNK1D*), CK1ε, and CK1γ2 (*CSNK2G2*), while no binding occurred to the control kinase NME2 or to GFP (Fig. 1a and Extended Data Fig. 1a,b). In kinase assays, besides DDX3X, also DDX6, -20, -39A, -41, -50 and -56 enhanced CK1 *in vitro* kinase activity of isoforms α, ε, δ, and γ2, but with different preferences (Fig. 1a and Extended Data Fig. 1c). Purified DDX proteins on their own showed no kinase activity towards peptide substrate, ruling out contaminating CK1 kinases.

**Fig. 1.**
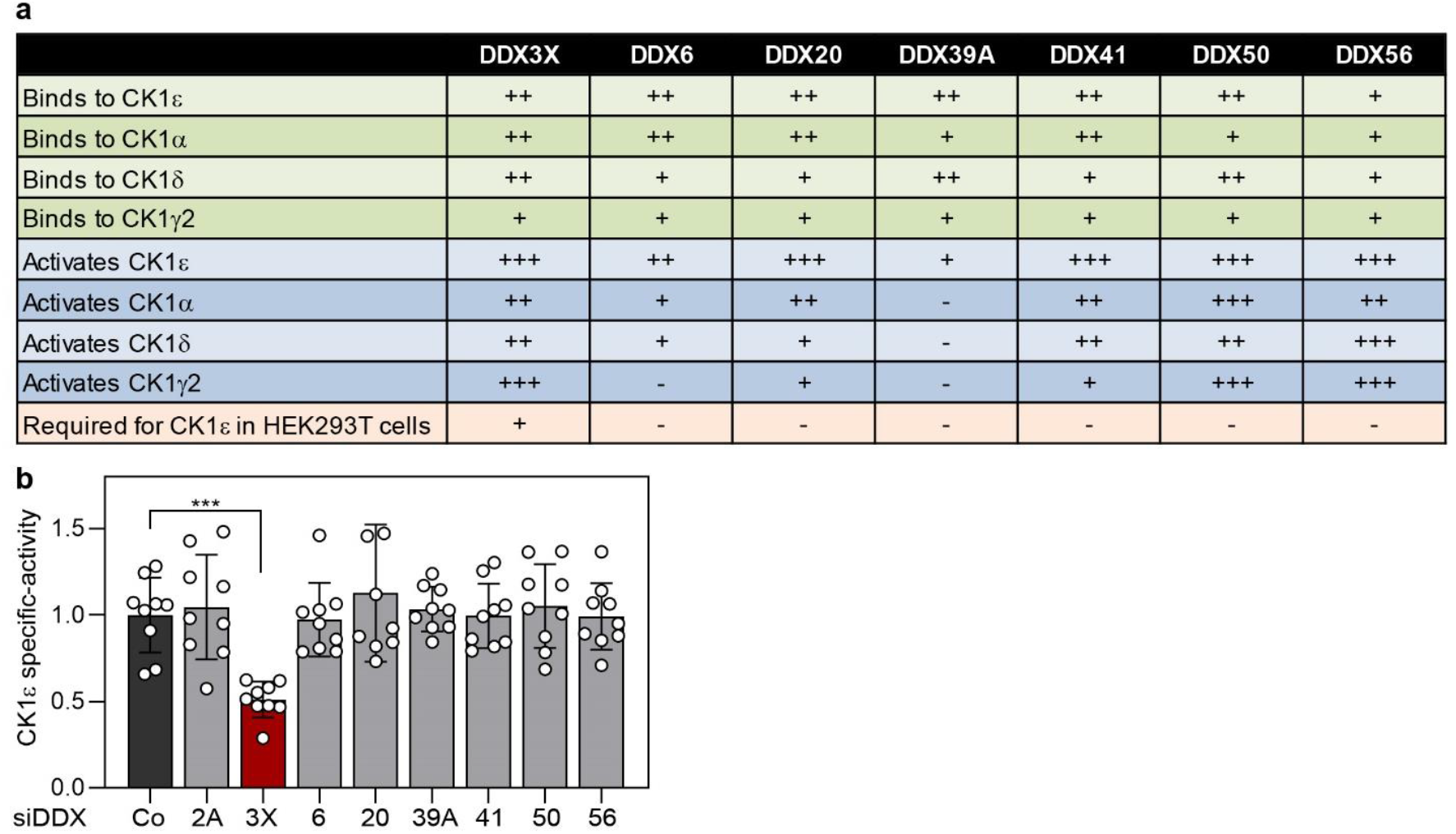
Latent Wnt/CK1ε -stimulating activity of DDX proteins revealed by chimeras. **a**, Summary of analyses for DDX-CK1 interactions from Extended Data Fig. 1 and Fig.1b. **b**, Kinase activity of CK1ε immunopurified from HEK293T cells and normalized to CK1ε protein levels (specific-activity) after knockdown of the indicated DDX protein. N = 3 (three independent experiments) and n = 9 (three biological replicates per independent experiment). Error bars are displayed as mean ± SD, ***: p < 0.001.

To test whether DDX proteins are not only sufficient *in vitro* but also required for full CK1ε activity in cells, we carried out kinase assays with CK1ε immunoisolated from HEK293T cells after *DDX* siRNA treatment. Importantly, to avoid confounding indirect or unspecific effects on kinase protein levels upon *DDX* siRNA treatment, we determined the *specific-*activity of the kinase, i.e. the kinase activity exhibited per molecule of kinase, by normalizing CK1ε activity in initial rate reactions to CK1ε protein levels determined by ELISA from the very same IP. Thus, even if cell growth or CK1ε expression was compromised due to *siDDX* affecting transcription or translation, our measurements control for such effects. While si*DDX3X* reduced CK1ε kinase specific-activity as reported^3^, siRNAs against *-6, -20*, -*39A, -41*, -*50* and -*56* had no effect (Fig. 1a,b and Extended Data Fig. 1d,e). Even siRNA knockdown of *DDX2A (EIF4A-1)*, the most abundant cellular DDX and “godfather” of DEAD-box proteins^9^, which caused a massive cell growth arrest (Extended Data Fig. 2c), had no effect on CK1ε specific-activity, attesting to the specificity of the si*DDX3X* effect and the validity of our specific-activity kinase assay. We conclude that while *in vitro* several DDX proteins can bind and stimulate CK1ε, *in vivo* the requirement for full CK1ε kinase activity is highly specific.

### Full CK2 kinase activity requires multiple DDX proteins in vertebrate cells

The *in vitro* activity of multiple DDX helicases suggested that their ability for kinase stimulation might not be limited to CK1. To address this possibility, we conducted several molecular screens, details of which will be reported elsewhere, that suggested casein kinase 2 (CK2) as a candidate susceptible to DDX stimulation. CK2 is a ubiquitous, pleiotropic and constitutively active protein kinase unrelated to CK1. CK2 plays key roles in cell proliferation, transcription, and signalling, and is essential in mice^10^. To study whether CK2 is physiologically activated by a DDX protein, we searched the CK2 interactome reported in HeLa cells^11^ and retrieved DDX24, -41, and -54 as potential regulators. CoIP of endogenous CK2α1 confirmed interaction with all three DDX proteins, but not with DDX56, indicating that the interaction is specific (Fig. 2a). Furthermore, we confirmed activation of CK2α2 *in vitro* by DDX41^169-569^ (Fig. 2b). To test whether these DDX proteins are required to stimulate CK2 in cells, we carried out kinase assays with CK2α1 immunoisolated from HeLa cells after *DDX* siRNA treatment. Importantly, combined DDX24/41/54 siRNA treatment of all three DDX proteins in HeLa cells reduced CK2α1 specific-activity without affecting cell growth (Fig. 2c and Extended Data Fig. 2a-c). Conversely, siRNA knockdown of *DDX2A*, although strongly affecting cell growth, had no effect on CK2 kinase specific-activity (Extended Data Fig. 2c-d). Likewise, combined DDX24/41/54 siRNA treatment reduced phosphorylation of a well-established CK2 phosphorylation target, HSP90 Ser226^12^, relative to total HSP90 protein (Fig. 2d,e). Thus, DDX24, -41, and -54 interact with CK2α1 and their combined action is required for full CK2α1 specific-activity.

**Fig. 2.**
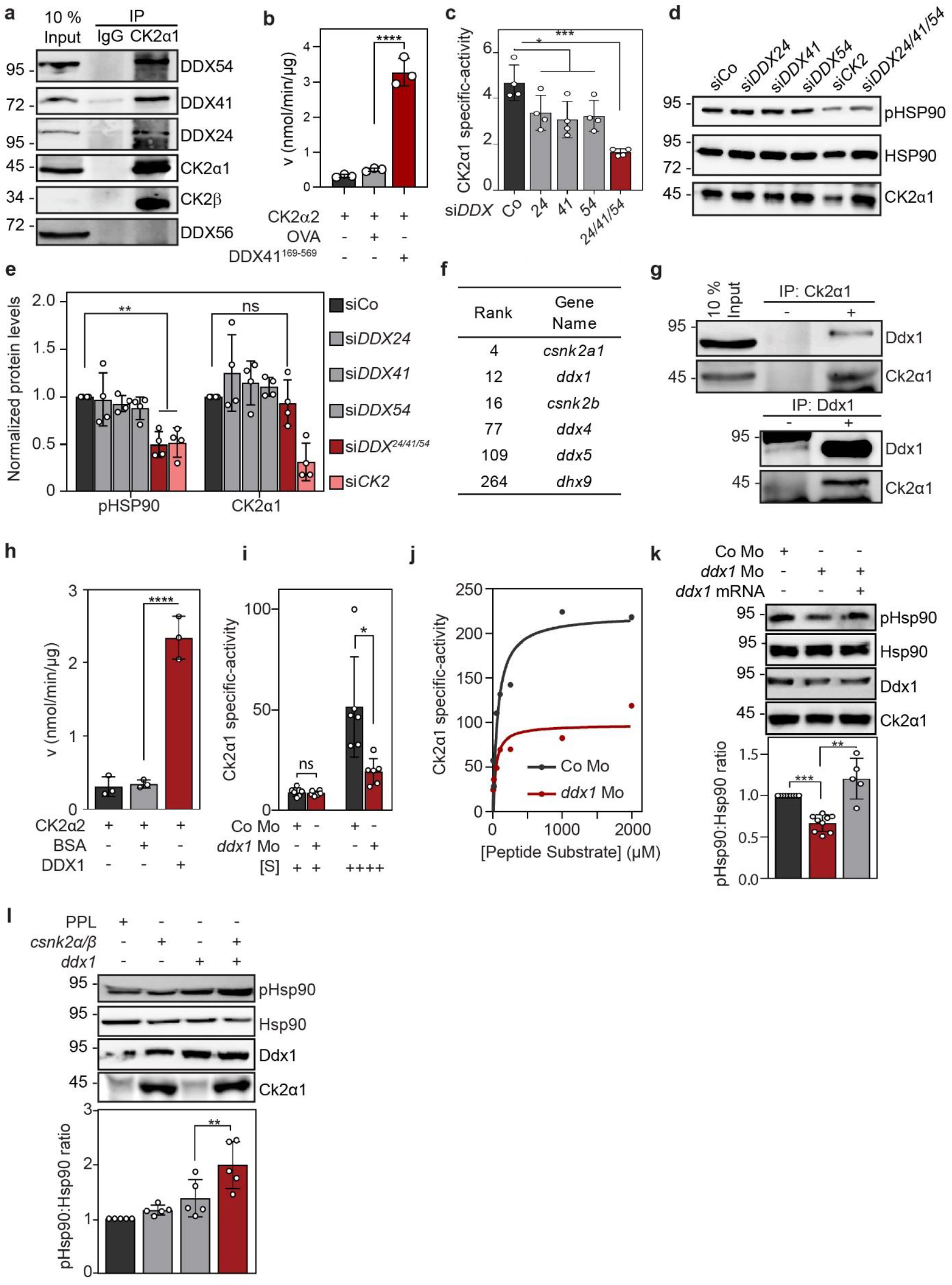
DDX1, 24, 41, and-54 stimulate CK2α1 in mammalian cells and frog embryos. **a**, Western blot of co-immunoprecipitation (CoIP) experiment from HeLa cells between endogenous CK2α1 and the indicated proteins. DDX56 and CK2β were included as negative and positive control respectively. **b**, *In vitro* kinase assay using recombinant CK2α2 and DDX41^169-569^ or ovalbumin (control). [Peptide substrate] = 500 µM. n = 3. **c**, Specific CK2α1 activity as determined from immunopurified and normalized CK2α1 protein from HeLa cells after siRNA knockdown of either DDX24/41/54 alone, in combination or control siRNA (siCo) treatment. n = 4 biological replicates. **d**, Western blot analysis of phospho-HSP90 (pHSP90) in HeLa cells after siRNA knockdown of DDX24/41/54 either alone or in combination. Knockdown of CK2 subunits α1, α2, and β was included as positive control (siCK2). **e**, Quantification of pHSP90 normalized to total HSP90 in four independent Western blot experiments from (**d**). **f**, Selected hits from a Ck2α1 interactome analysis by mass-spectrometry of proteins CoIP with Ck2α1 in *X. laevis* embryos (stage 18) showing ranks of selected hits. **g**, Western blot of CoIP from *X. tropicalis* embryo extracts (stage 18) between endogenous Ddx1 and Ck2α1 and in both directions (top, bottom). **h**, *In vitro* kinase assay of recombinant CK2α2 with DDX1 or BSA (control). n = 3. **i**, Specific Ck2αactivity as determined from immunopurified and normalized Ck2αprotein from *X. tropicalis* embryos injected with either Co or *ddx1* Mo. Assays were carried out at low (2 µM, +) or high (1.0 mM, ++) peptide substrate concentration. n = 6 biological replicates. **j**, Ck2α1 specific-activity at increasing peptide substrate concentrations with Ck2αimmunopurified from *X. tropicalis*, injected with either Co or *ddx1* Mo. **k**, Top, western blot of pHsp90 in *X. tropicalis* embryos (stage 18) that were injected at two-to four-cell stage with *ddx1* Mo. Where indicated, non-targetable *ddx1* mRNA was co-injected. Bottom, quantification of western blot results: n= 9 (Co and *ddx1* Mo), n= 5 (*ddx1* Mo + *ddx1* mRNA) biological replicates. **l**, Top, western blot of *X. laevis* embryos that were injected with either *csnk2α/β* or *ddx1* mRNA alone or in combination. Bottom, quantification of western blot results: n = 5 biological replicates. Data shown are representatives of at least two independent experiments and where indicated are displayed as means ± SD with one-way ANOVA (**b, c, e, h, k, l**) or paired t-test (**i**). * p < 0.05, ** p < 0.01, *** p < 0.001

To corroborate that CK2α1 is physiologically DDX-regulated also *in vivo*, we analysed *Xenopus tropicalis* embryos and used CoIP-mass-spectrometry to identify the Ck2α1 interactome. Among the Ck2α1-binding proteins identified, Ddx1 was a top hit (Fig. 2f, Extended Data Table 1). *ddx1* is an ubiquitously expressed, essential gene with roles in splicing, RNA processing, and – transport^13^. CoIP of the endogenous proteins (Fig. 2g) confirmed the interaction of Ddx1 and Ck2α1 in *Xenopus* embryos. *In vitro* kinase assays showed that recombinant DDX1 potently stimulates CK2α2 kinase activity (Fig. 2h).

To analyse its requirement, we depleted Ddx1 protein levels in *Xenopus* embryos by injecting an antisense morpholino (Mo). We validated Mo specificity by confirming that the Mo indeed reduced Ddx1 protein levels (Extended Data Fig. 2e). To monitor endogenous kinase activity, we carried out initial rate *in vitro* kinase assays with Ck2α1 immunopurified from control embryos or *ddx1* morphants. *ddx1* Mo injection reduced Ck2α1 specific-activity, which, interestingly, manifested predominantly at high peptide-substrate concentration (Fig. 2i-j, see Extended Data Fig. 2f for assay linearity). Moreover, *ddx1* Mo reduced Hsp90 phosphorylation relative to total Hsp90 protein (Fig. 2k and Extended Data Fig. 2g). Co-injection of *ddx1* mRNA rescued Hsp90 phosphorylation in Ddx1 morphants and Ddx1 and Ck2α1 overexpression synergistically induced Hsp90 phosphorylation (Fig. 2k,l).

Taken together, we conclude that i) multiple DDX proteins stimulate cellular CK2 *in vivo*, ii) that the requirement of DDX proteins for full kinase specific-activity is evolutionary conserved between frog and human, iii) that the requirement of DDX proteins for full kinase specific-activity is not limited to DDX3X and CK1ε, and iv) that kinase stimulation by DDX manifests at high substrate concentration.

### Kinase stimulation by DDX3X requires RNA binding-but not catalytic motifs

The ability of multiple DDX proteins to stimulate both CK1 and CK2 suggests that the conserved core domain mediates kinase stimulation and raises the question, which motifs are involved. To study this question, we produced recombinant proteins in *E. coli* for DDX core domains, CK1ε ^1- 294^ (lacking the autoinhibitory tail), and CK2α2. We selected DDX3X and a panel of six unrelated helicases including DDX5, -6, -17, -19A, -27, and -56, that we arbitrarily chose either because we had tested them previously^3^ or because they produced well as recombinant proteins (Extended Data Fig. 3a,b) and that were functional and bound RNA (Extended Data Fig. 3c). Initial rate reactions showed that to varying degree, all DDX core domains but not BSA or GFP stimulated CK1ε ^1-294^ and CK2α2 kinase activity towards kinase-specific peptide substrate and casein protein (Extended Data Fig. 3d-g).

The DDX core domain harbours the catalytic activity, as well as ATP/ADP- and RNA binding motifs. To test if either ADP binding or ATPase activity is required for protein kinase stimulation, we produced DDX3X core domain proteins harbouring oncogenic mutations^14^ (Extended Data Fig. 4a,b) that inhibited ADP binding (R528H) (Extended Data Fig. 4c), or that reduced or abolished ATPase activity (R276K, D506Y, R528H, R534H) (Extended Data Fig. 4d). We also tested a well-characterized ATPase-dead form, DADA (D346A/D349A)^15^ that binds ADP but fails to hydrolyse ATP (Extended Data Fig. 4c,d). However, none of these mutants was much impaired in stimulating CK1ε kinase activity *in vitro* (Extended Data Fig. 4e). Likewise, the saturation kinetics of the DADA mutant, devoid of ATPase activity, was indistinguishable from that of wild type DDX3X (Extended Data Fig. 4f). Thus, DDX3X ATPase activity is not required for casein kinase stimulation.

To identify motifs required for kinase-stimulation, guided by the available crystal structures^16,17^ we conducted alanine-scanning mutant analyses of amino acids located on the DDX3X surface (Fig. 3a). In a preliminary mutational analysis, we found that RecA-like domains 1 and 2 (D1 / D2) both contribute to kinase activation and we therefore created compound mutants. We iteratively generated and tested single-to-quintuple mutants and identified in both domains a patch of mostly positively charged amino acids required for CK2α2 and CK1ε ^1−294^ *in vitro* kinase stimulation (Fig. 3a-c). We further characterized representative mutants active-(#3, 8) or inactive (#16 ; #17) in both, *in vitro* kinase stimulation (Fig. 3 b,c) and Wnt signalling (Fig. 3d,e). Overall folding of all four mutants was unaffected as judged either by thermal stability assays (Fig. 3f) or by their ability to bind a fluorescent ADP derivative (2’/3’-O-N-Methyl-anthraniloyl (mant) ADP) (Fig. 3g). However, the two kinase-inactive DDX3X mutants #16 and #17 showed strongly reduced RNA binding ability (Fig. 3h,i). Projection of the mutated residues onto the DDX3X crystal structure reveals two basic patches in the D1 (located in an unstructured loop) and D2 domain (Fig. 3j), of which two residues in D2 (K451, G473) are already known to contact RNA^17^. These results suggest that RNA binding and kinase stimulation in DDX3X may be mutually exclusive (Fig. 3k). Consistently, we found that poly-uridine RNA inhibits the ability of all tested DDX proteins to stimulate CK2α2 kinase activity, while RNA had little effect on the kinase alone (Fig 3l). We therefore conclude that RNA binding and kinase stimulation in DDX3X require common motifs, supporting that kinase regulation involves the RNA client-recognition interface in DDX3X.

**Fig. 3.**
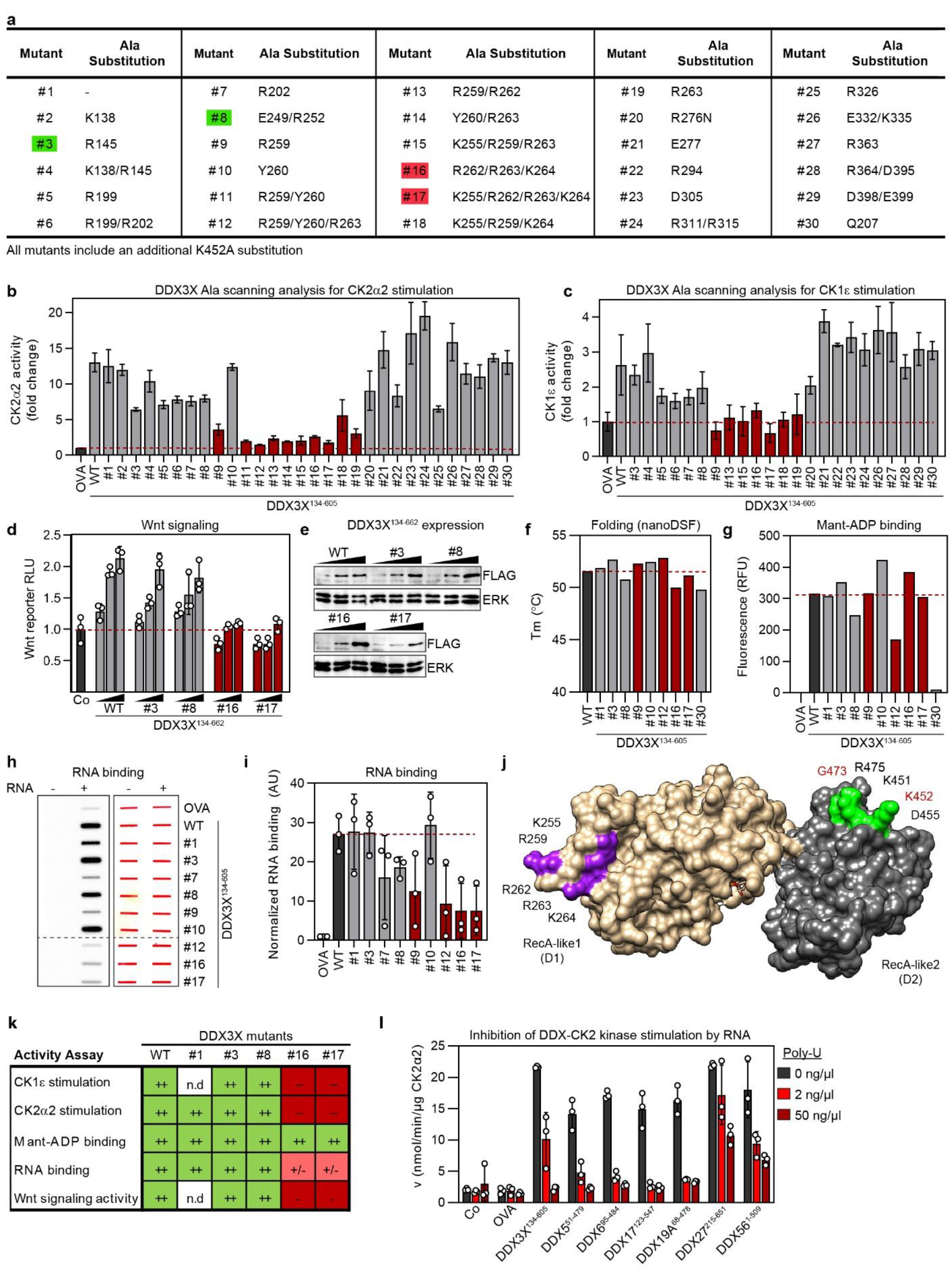
DDX3X Mutants fail to stimulate CK2α2 and CK1ε ^1-294^ kinase activity. **a**, Table of DDX3X^134-605^ alanine scanning mutants analysed. All mutants harboured an additional K452A substitution in D2. Key mutants shown in Fig. 3m are highlighted; kinase stimulation impaired (red), unaffected (green). **b**, *In vitro* kinase assay with recombinant CK2α2 and DDX3X^134-605^ mutant proteins. **c**, *In vitro* kinase assay with recombinant CK1ε ^1−294^and DDX3X^134-605^ mutant proteins. **d**, Wnt-reporter (TopFlash) assay in HEK293T cells after transfection with increasing doses of the indicated DDX3X^134-662^ mutants, i.e. containing the CTD required for Wnt activation. In all assays, cells were cotransfected with *siDDX3X* and treated with Wnt3a. Red bars indicate mutants deficient in kinase stimulation. **e**, Western blot of (**d**) confirming equal transfection amounts of the DDX3X^134-662^ mutants. **f**, Thermal stability analysis by nanoDSF for selected DDX3X^134-605^ mutants. Red bars indicate mutants deficient in kinase stimulation. **g**, mant-ADP binding analysis for selected DDX3X^134-605^ mutants. Red bars indicate mutants deficient in kinase stimulation. Mutant #30 (Q207A) was specifically designed to abolish ADP binding. **h**, RNA (PolyA^-^) binding analysis (left panel) for selected DDX3X^134-605^ mutants by filterblot. Ponceau staining (right panel) shows equal loading of the proteins. **i**, Quantification of RNA binding in (**h**). Red bars indicate mutants deficient in kinase stimulation. **j**, DDX3X X-ray crystal structure with surface rendering (PDB ID: 5E7J^16^). The RecA-like domains are shown in beige (D1) and grey (D2). Alanine-substitutions of amino acids that abolish CK1ε and CK2α2 kinase stimulation are in purple (D1) and green (D2). Note that amino acids K451 and G473 in D2 are part of the known RNA binding interface^17^. **k**, Summary of mutant analyses by alanine-scanning of DDX3X. n.d. = not determined. (++), (+/-), (-) indicate activity relative to wt DDX3X core domain or DDX3^134-662^ respectively. **l**, *In vitro* CK2α2 kinase assay in the presence of DDX core domain proteins and increasing concentrations of synthetic RNA (poly-uridine). Data shown are representatives of at least two independent experiments (with n = 3) and where indicated are displayed as means ± SD

### DDX proteins act as nucleotide exchange factors for CK2 to relieve substrate inhibition

The ability of DDX proteins to stimulate multiple protein kinases notably at high substrate concentration suggests a universal underlying kinetic mechanism. To elucidate the kinetic mechanism, we carried out a comprehensive steady-state enzyme kinetics analysis and first determined the catalytic mechanism of CK1ε ^1-294^ and CK2α2 in the absence of DDX. We carried out two-substrate steady-state kinetic analysis and measured initial reaction velocities at varying concentrations of peptide substrate, mutant peptide, ATP, and ADP. We derived the corresponding velocity equation for a general sequential bi-bi mechanism (Fig. 4a) and using a maximum-likelihood approach determined its kinetic parameters by globally fitting this equation simultaneously to all velocity data for a given enzyme. The best fits indicate an ordered mode of substrate-cosubstrate binding for CK1ε ^1-294^ (ATP first, peptide substrate second) and a random mode for CK2α2 (Fig. 4a,b and Extended Data Fig. 5-6 for individual [ATP] panels). The kinetic parameters for both enzymes were well constrained by the experimental data (Extended Data Fig. 5b, 6b); hence, the model can be used to make quantitative predictions. Interestingly, when increasing the peptide substrate concentrations, the velocity curves rise to a maximum and then decline (Fig. 4b), known as substrate inhibition, a phenomenon whereby enzymes are inhibited at high concentrations of their own substrate. This substrate inhibition is predicted by our model to be due to ordered product release for both CK1ε ^1-294^ and CK2α2, with phosphorylated peptide dissociating prior to ADP (Fig. 4a, Supplementary Information). As a result, a dead-end complex, with both ADP and peptide substrate bound to the kinase, becomes prevalent at high peptide substrate concentrations. In bi-substrate enzyme reactions such e.g. protein kinases, substrate inhibition by unproductive binding at the catalytic site is the rule^18^ and in protein kinases, ADP release is commonly rate-limiting^19^ and may cause substrate inhibition.

**Fig. 4.**
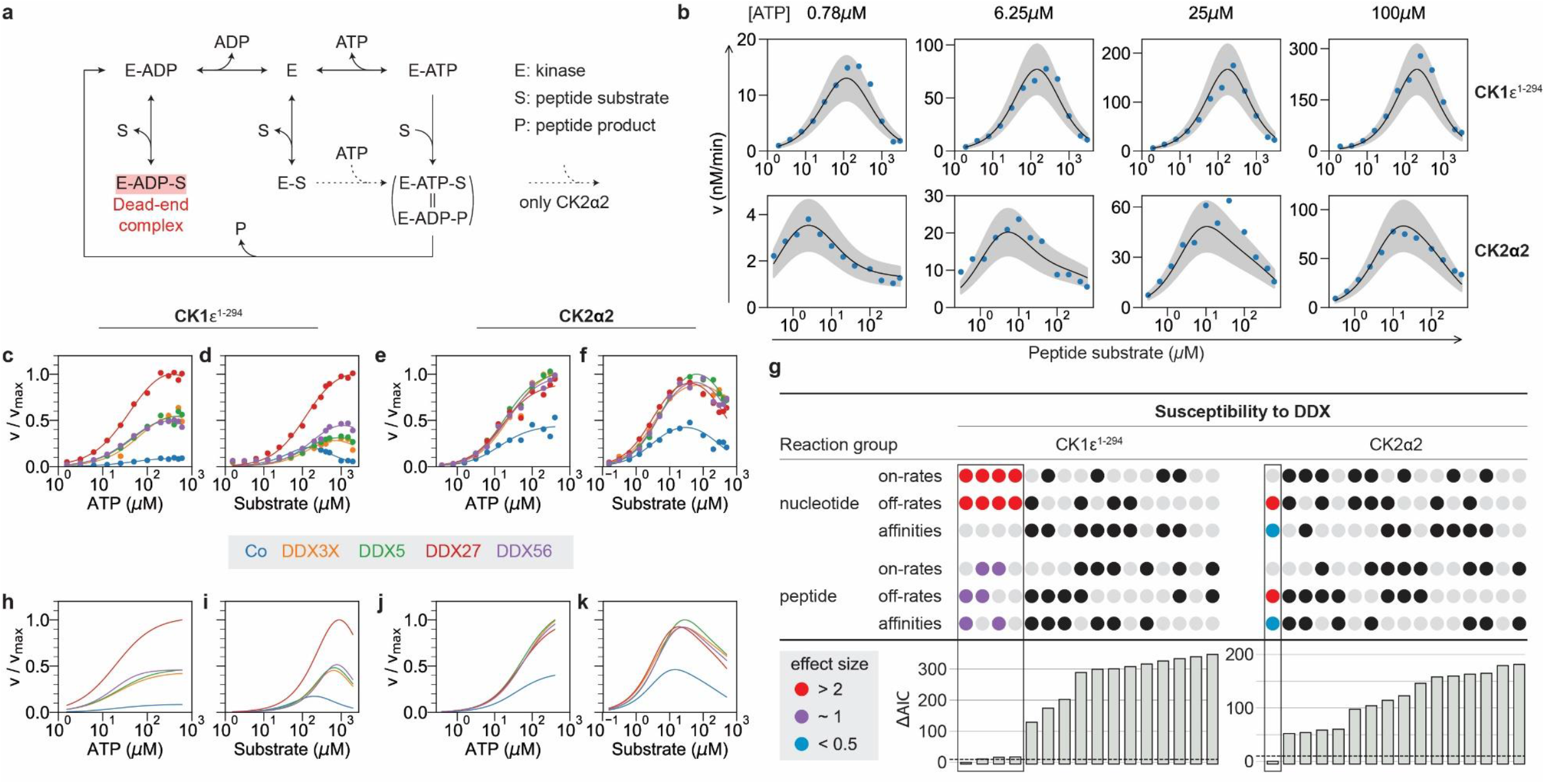
DDX proteins accelerate the nucleotide off-rate in CK1ε ^1-294^ and CK2α2. **a**, Scheme for a sequential bi-bi reaction mechanism of kinases, allowing for ordered (CK1ε ^1-294^) or random (CK2α2) binding of substrates (peptide, ATP), while ordered product release (phospho-peptide preceding ADP) results in a dead-end complex underlying substrate inhibition. **b**, Initial rate measurements for CK1ε ^1-294^ (top) and CK2α2 (bottom) upon systematic variation of the concentrations of peptide substrate and ATP. Data were fitted with ordered- and random substrate binding models for CK1ε ^1-294^ and CK2α2, respectively (see **a**). Blue dots, experimental data; black curves, best model fit; grey shades, ±1 SD error bands. **c-f**, Initial rate measurements in the presence or absence of DDX proteins as indicated. **c**, for CK1ε ^1-294^ at varying concentrations of ATP; **d**, for CK1ε ^1-294^ at varying concentrations of peptide substrate; **e**, for CK2α2 at varying concentrations of ATP and [peptide] = 20 µM; **f**, for CK2α2 at varying concentrations of peptide substrate. **g**, Prediction of the mechanism of DDX action by statistical model selection with substrate inhibition by dead-end complex formation. DDX proteins were allowed to affect on- or off-rates for adenine nucleotides (ATP/ADP) or peptide, either as single parameters or in any combination, constrained by equal effect sizes for a given pair of on- and off-rates when both were allowed to change upon DDX binding (upper panels). The best fits of each model to the measured velocity data were ranked by the Akaike information criterion (AIC, lower panel). The effect sizes for DDX27 are indicated for the models close to or passing the standard selection threshold (ΔAIC < 10). **h-k**, Fit of the best models for CK1ε ^1-294^ (**h-i**) and CK2α2 (**j**-**k**), having the lowest ΔAIC, to the corresponding reaction-velocity measurements shown in **c-f**. n = 1 (**b**) and n = 2 (**c**-**f**).

Next, to identify the mechanisms of action of DDX proteins, we measured initial rate reactions for CK1ε ^1-294^ and CK2α2 at varying concentrations of peptide and ATP in the presence of DDX3X, - 5, -27, and -56 (Fig. 4c-f). DDX proteins caused no major change in the apparent Michaelis-Menten constant K_m_ for the peptide substrate. However, they strongly increased, to different extent in both kinases, the apparent inhibition constant for substrate inhibition K_i_ and the maximal reaction velocity V_max_ (Extended Data Table 2). In CK1ε ^1-294^, DDX proteins mostly relieved substrate inhibition, increasing K_i_ greater than 10-fold (Figure 4d; kinetic constants in Extended Data Table 2). In CK2α2, there was a smaller (up to 4-fold) increase in K_i_ and, additionally, a 2-fold increase in V_max_ (Figure 4f; kinetic constants in Extended Data Table 2). Release from substrate inhibition in CK2α2 is consistent with the observation in *Xenopus* that deficiency of Ddx1 manifested predominantly at high substrate concentration (Fig. 2i,j). Varying ATP concentrations, DDX proteins had the same effect on both kinases, increasing V_max_ (Fig. 4c,e; kinetic constants in Extended Data Table 2). In sum, this analysis indicates effects of DDX proteins on substrate inhibition and V_max_. To infer the underlying mechanisms, we used the above models and asked which elementary rate parameters for CK1ε ^1-294^ and CK2α2 change when fitting the models to the reaction velocity measurements in the presence of DDX.

Assuming that phosphoryl-transfer is not rate-limiting^19^, we tested systematically whether DDX proteins affected on- or off-rates of nucleotides (ATP/ADP) or peptide/phospho-peptide, either as a single parameter or in all possible combinations. We fitted the resulting 16 models to the experimental data, ranked DDX susceptibility of the models by the Akaike Information Criterion (AIC)^20^, and selected models with a difference of ΔAIC < 10 (Fig. 4g)^20,21^. For CK1ε ^1-294^, four very similar models were selected that all show strongly accelerated nucleotide on- and off-rates; the effects on peptide binding or release were small (Fig. 4g; fits of best model to data shown in Fig. 4h-i; see Extended Data Fig. 7a,b for individual DDX panels). For CK2α2, the only model selected shows enhanced nucleotide release by DDX proteins as well as an acceleration of peptide release by a similar magnitude (Fig. 4g,j,k; see Extended Data Fig. 7c,d for individual DDX panels). Importantly, the common feature of models selected for CK1ε ^1-294^ and CK2α2 is that DDX proteins are predicted to enhance the nucleotide off-rate, i.e. to act as nucleotide exchange factors (NEF). We also modelled the kinase data assuming inhibition by allosteric binding of the substrate, rather than by forming a dead-end complex. The best fitting models also support accelerated nucleotide release by DDX proteins (Extended Data Fig. 8). Thus, the conclusion based on the above enzyme kinetics data that DDX proteins act as NEF is robust.

To corroborate this conclusion, we confronted the selected models for DDX action with experimental data not used for training. First, we monitored the effect of DDX proteins on the inhibition of CK1ε ^1-294^ and CK2α2 by varying ADP. DDX proteins did not relieve product inhibition by ADP (Fig. 5a,b). This was correctly predicted by the models (Fig. 5c,d) and is due to DDX proteins increasing the on- and/or off-rate of both ADP <u>and</u> ATP. Next, we tested whether DDX proteins relieve competitive inhibition of the kinases by non-phosphorylatable mutant peptides carrying Ser to Ala substitution, which was the case for both kinases (Fig. 5e,f). The models reproduced the observed effects, at least qualitatively (Fig. 5g,h). Thus, the acceleration of nucleotide exchange by DDX proteins (and, potentially peptide release) does not only relieve substrate inhibition but counters also the effects of competitive substrate inhibitors. Lastly, we studied the dose-dependence of the DDX effect. All tested DDX proteins increased the reaction velocities of the kinases in a dose-dependent manner with EC_50_ between ∼0.1-1.0 μM and Hill coefficients n_H_ of ∼1.0 (Fig. 5i,j, Extended Data Table 2), whereas none of the control proteins (GFP, BSA, γ-globulin) did, even at much higher concentrations (Extended Data Fig. 7e). The DDX dose-effects were readily reproduced by the selected models (Fig. 5k,l; see Extended Data Fig. 7f for individual DDX panels).

**Fig. 5.**
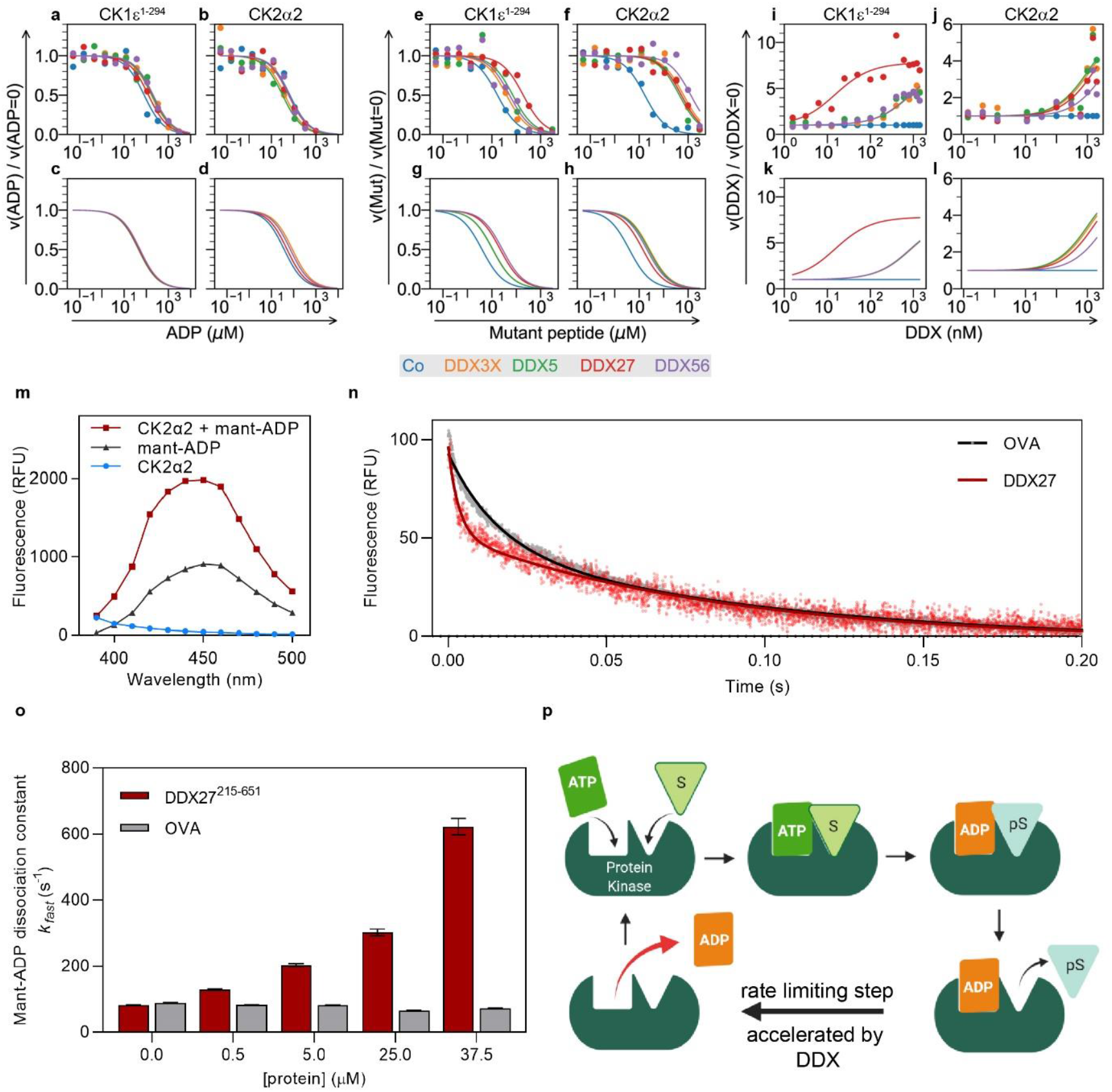
DDX proteins function as nucleotide exchange factor for CK2α2. **a-d**, Addition of DDX proteins does not relieve product inhibition by ADP neither for CK1ε ^1-294^(**a**) nor for CK2α2 (**b**, [peptide] = 40 µM), as predicted by the selected best models for CK1ε ^1-294^(**c**) and CK2α2 (**d**). **e-h**, Addition of DDX proteins relieves competitive inhibition by non-phosphorylatable mutant peptide substrates, for both CK1ε ^1-294^ (**e**, [peptide] = 150 µM) and CK2α2 (**f**, [peptide] = 150 µM), as qualitatively predicted by the selected best models for CK1ε ^1-294^ (**g**) and CK2α2 (**h**), respectively. **i-l**, Dose-responses of reaction velocity against concentration of different DDX proteins measured for CK1ε ^1-294^ (**i**) and CK2α2 (**j**) are reproduced by the selected best models for CK1ε ^1-294^ (**k**) and CK2α2 (**l**), respectively. Experimental data show one representative experiment of n = 2 (**a, b, e, f, i, j**). **m**, Fluorescence spectra of mant-ADP upon excitation at 290 nm, in presence and absence of CK2α2, representing the FRET signal employed in stopped-flow analyses in (**n, o**). **n**, Representative stopped-flow analysis of the time-resolved dissociation of mant-ADP from CK2α2 in presence of ovalbumin and DDX27^215-651^ (25 µM final concentration). CK2α2 (1 μM) was pre-equilibrated with mant-ADP (20 μM) in one syringe and rapidly mixed at 1:1 ratio with a solution containing ADP (5 mM) and 50 µM ovalbumin (OVA, negative control) or DDX27^215- 651^. Raw data and fitted curves are superimposed. **o**, Mant-ADP dissociation constants of CK2α2 in presence of the indicated final concentrations of DDX27^215-651^ or ovalbumin from stopped-flow measurements as in (**n**). Data were fitted to a two-phase decay exponential function to obtain dissociation rate constants. Error bars indicate standard error of the fit. **p**, Model for DDX proteins acting as nucleotide exchange factors for protein kinases. In protein kinases, ADP release is commonly the rate-limiting step in the reaction. Rebinding of substrate may occur, leading to a [ADP-kinase-substrate] dead-end complex (not shown). DDX binding to protein kinase accelerates ADP release and reduces dead-end complex formation, thereby increasing reaction-rate, notably at high substrate concentrations. Created with Biorender.com.

To analyse directly whether DDX proteins accelerate nucleotide exchange, we measured the dissociation kinetics of mant-ADP from CK2α2 by stopped-flow spectroscopy, the ‘gold-standard’ for determining NEF activity^22^. In steady-state fluorescence measurements, we first established that binding to CK2α2 increased mant-ADP fluorescence due to resonance energy transfer (FRET) (Fig. 5m). Next, we monitored the FRET signal from the CK2α2-mant-ADP interaction in stopped-flow spectroscopy measurements. The dissociation rate constant for the mant-ADP was determined by preincubating CK2α2 with mant-ADP and then rapidly mixing with a large excess of unlabelled ADP, which displaces mant-ADP from the kinase. The concomitant time-resolved decrease in fluorescence was recorded and the dissociation rate constant for CK2α2 determined as 85 ± 4s^-1^ (8.1 ms half-life) (Fig. 5n,o). This value is in the order of nucleotide-dissociation rates obtained for protein kinase A^22^. We then tested the effect of DDX27^215-651^ on the ADP dissociation CK2α2 because its own ADP binding, which can interfere with measurements, was low compared to other DDX proteins. Importantly, DDX27^215-651^ accelerated mant-ADP dissociation from CK2α2 in a dose-dependent manner up to ∼7-fold, while the control protein ovalbumin had no effect (Fig. 5n,o, Extended Data Table 2). We conclude that DDX27^215-651^ accelerates the ADP off-rate of CK2α2 and hence exhibits NEF activity.

## Discussion

In this study, we provide evidence for a role DDX proteins as kinetic modifiers of protein kinase activity and we reveal NEF activity in DDX proteins towards CK2α2. This notion is highly unexpected and has potentially wide-ranging implications. The data also raise the possibility of functional coupling between DDX and protein kinase activity and thus links between cellular signaling and RNA processing. Concerning specificity, while *in vitro* several DDX proteins can interact with and stimulate CK1ε and CK2α2, *in vivo* the requirement for full kinase activity by their cognate DDX partners is selective, suggesting specificity determinants residing in the non-conserved helicase domains that remain to be elucidated.

In protein kinases, ADP release is commonly rate-limiting and hence the compelling proposition was made that cells may leverage nucleotide exchange to control protein kinase activity^23^, but such NEFs have remained elusive. The results from steady-state kinetics, global modelling, prediction-validation, and stopped-flow spectroscopy converge on the unexpected conclusion that DDX proteins act as NEFs to stimulate CK2α2 (Fig. 5p). Kinetic modelling predicts this also to be the case for CK1, but low FRET signals prevented us from confirming the prediction by stopped-flow analysis. Our data support a direct effect of DDX on casein kinase stimulation also *in vivo* as indicated i) by the requirement for full specific-activity of CK1ε for DDX3X^3^ and of CK2α2 for DDX1/24/41/54 in human cells and in frog embryos, ii) the fact that the DDX stimulation towards CK2α2 *in vivo* manifests specifically at high [S] when [E-ADP-S] dead-end complexes become rate limiting.

Although at first glance their NEF activity appears surprising, DDX helicases are reminiscent of another NEF family, namely protein co-chaperones of the heat shock protein Hsp110 family^24^, while DDX proteins are chaperones for RNA folding. For example, like DDX3X, Hsp110 is a chaperone that does not require ATP hydrolysis for its NEF activity^25^. Like HSPs, DDX helicases are chaperones but they stabilize certain conformations in RNA instead of in proteins. We propose that DDXs can also stabilize an active kinase conformation that accelerates ADP release^26^. Supporting such a dual function, we find that basic amino acid residues required for kinase stimulation and RNA binding overlap in DDX3X, and that RNA prevents DDX from stimulating CK2α2 kinase activity. Finally, our simulations predict that for the CK2α2 reaction cycle, DDX proteins may not only accelerate release of ADP but also of peptide to speed up the overall reaction. While we did not directly analyze peptide release rate directly, this suggests multiple modes whereby DDX can modify kinase reaction kinetics.

There is a major difference between protein kinases and other NEF targets. Nucleotide dissociation of common NEF targets such as Hsp70 and GTPases is slow, with half-lives of protein-nucleotide complexes between seconds to hours. By comparison, ADP dissociation from protein kinases is fast, with protein-nucleotide complex half-lives in the centi-to millisecond range^22,23,27^. Hence, NEF activity of DDX proteins should be particularly important when substrate is in great excess over protein kinase leading to substrate inhibition, for example in protein condensates, where proteins can reach millimolar concentrations^28^. Protein kinases and DDX proteins are prominent regulators of such membraneless organelles^29,30^. We speculate that protein kinase reactions in condensates are a hot spot for DDXs to overcome substrate inhibition.

This study focused on selected DDX-casein kinase interactions, given 44 DDX proteins in humans^31^ whose function remains largely unknown, which could potentially interact with 518 protein kinases, there may be many unexplored kinase regulatory interactions.

## Methods

Methods are provided in the Supplementary Information

## Acknowledgements

We are grateful to L. Rohland and M. Mayer for help with stopped flow measurements. We thank C. Reinhard, N. Dirdjaja, and A. Wendlandt for expert technical assistance and J. Oakhill for advice and supply with P81 paper. We thank I. Dominguez, G. Maga, and B. Suter for providing constructs. We acknowledge G. Roth and Aska Pharmaceuticals Tokyo for generously providing hCG, and NXR (RRID: SCR_013731), Xenbase (RRID: SCR_004337), and EXRC for *Xenopus* resources. Expert technical support by the central animal laboratory of DKFZ is gratefully acknowledged.

## Funding

German Research Council (DFG) via the Collaborative Research Centre 1324 (TP B1 to C.N.) German Research Council (DFG) via the Collaborative Research Centre 1324 (TP B7 to I.S.) German Research Council (DFG) via the Collaborative Research Centre 873 (TP 11 to T.H.)

## Author contributions

E.F., A.H., A.S., and M.G. contributed equally to this work. E.F. conducted all kinase kinetics analyses and DDX characterization; A.H. validated CK2α2 interacting DDX proteins in human cell lines and *Xenopus*, and performed DDX3X mutant analyses. M.G. and T.H. conducted all kinetic modelling; A.S. produced recombinant proteins and carried out stopped flow analyses. G.S. contributed to initial recombinant protein production and performed DDX3X mutagenesis; C.M.C. conducted initial CK1 assays. D.P. and J.K. conducted proteomics analyses; G.S. and I.S. co-supervised A.S.. All authors analysed and discussed the data. C.N. conceived and coordinated the study and wrote the paper with contribution from T.H.

## Competing interests

Authors declare no competing interests.

## Data and materials availability

All data are available from the corresponding author upon request. The ProtoArray dataset is available in the Gene Expression Omnibus (GEO) under accession number GSE150859 [https://www.ncbi.nlm.nih.gov/geo/query/acc.cgi?acc=GSE150859]. The Mass-spectrometry dataset is available at ProteomeXchange under accession number PXD019405 [http://proteomecentral.proteomexchange.org/cgi/GetDataset?ID=PXD019405]. Custom computer programs for the mathematical modelling were coded in Python and will be made available upon request.

## Materials and Correspondence

Supplementary Information is available for this paper.

Correspondence and requests for materials should be addressed to Christof Niehrs

---

**Extended Data Fig.1.**
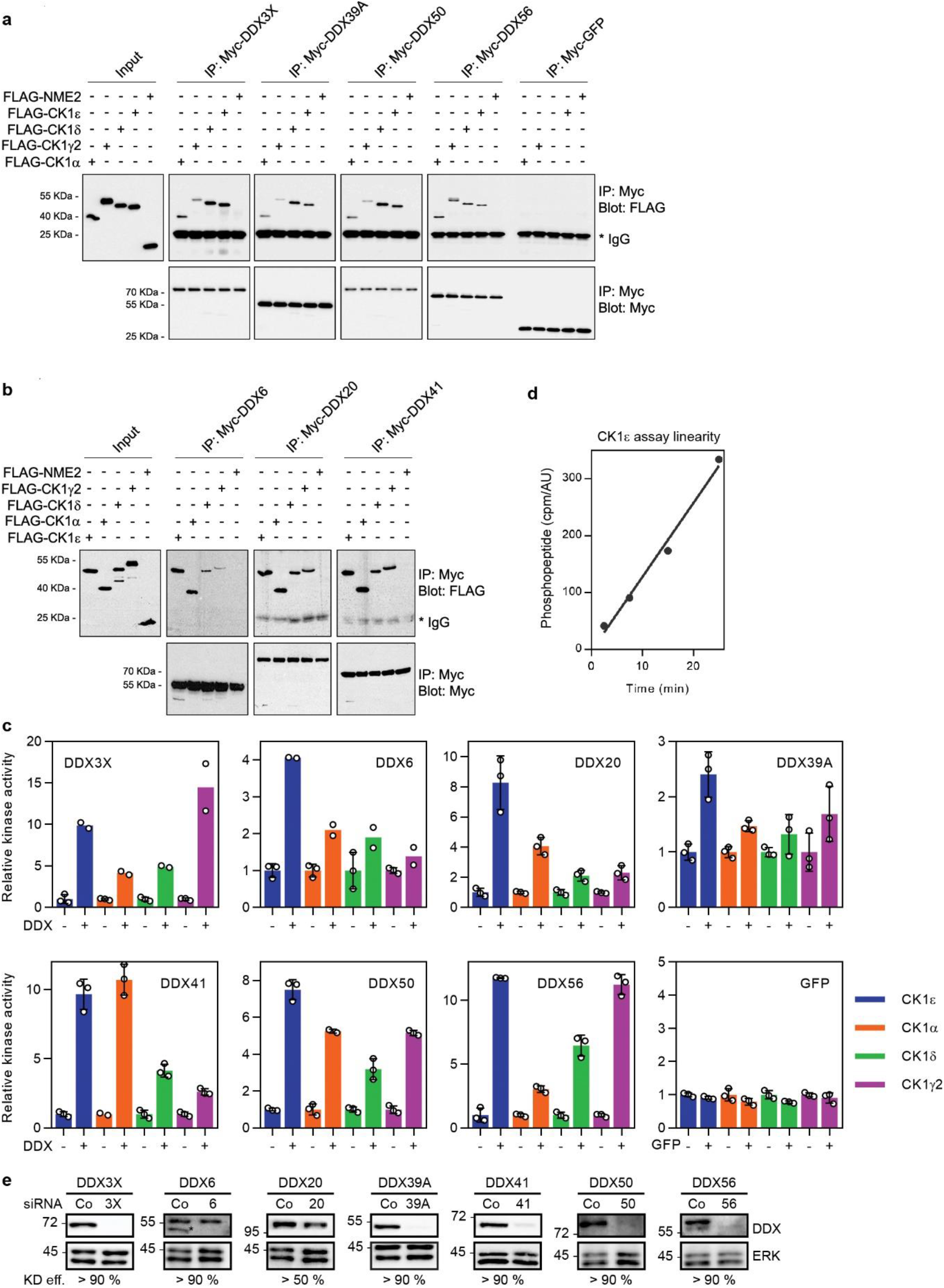
DDX proteins bind to and regulate CK1ε activity. **a-b**, *In vitro* binding assays with the indicated recombinant FLAG-tagged CK1 isoforms and immobilized Myc-tagged DDX proteins or GFP (control) from HEK293T cells. Western blotting was performed with anti-Myc and anti-FLAG antibody. FLAG-NME2 was included as a negative control. **c**, *In vitro* kinase assays with the indicated recombinant CK1 and specific peptide substrate in absence or presence of the indicated DDX protein. [Peptide substrate] = 50 µM. Data are displayed as means ± SD values. n = 2 – 3. **d**, *In vitro* kinase assay with CK1ε immunopurified from HEK293T cells to establish linearity for kinase assay in Fig. 1b. Phosphopeptide (cpm) was normalized to the kinase amount used (AU). N = 1. **e**, Western blot of HEK293T cells treated with the siRNA used in Fig. 1b, confirming knockdown of the corresponding DDX. The knockdown efficiency (“KD eff.”) is shown.

**Extended Data Fig.2.**
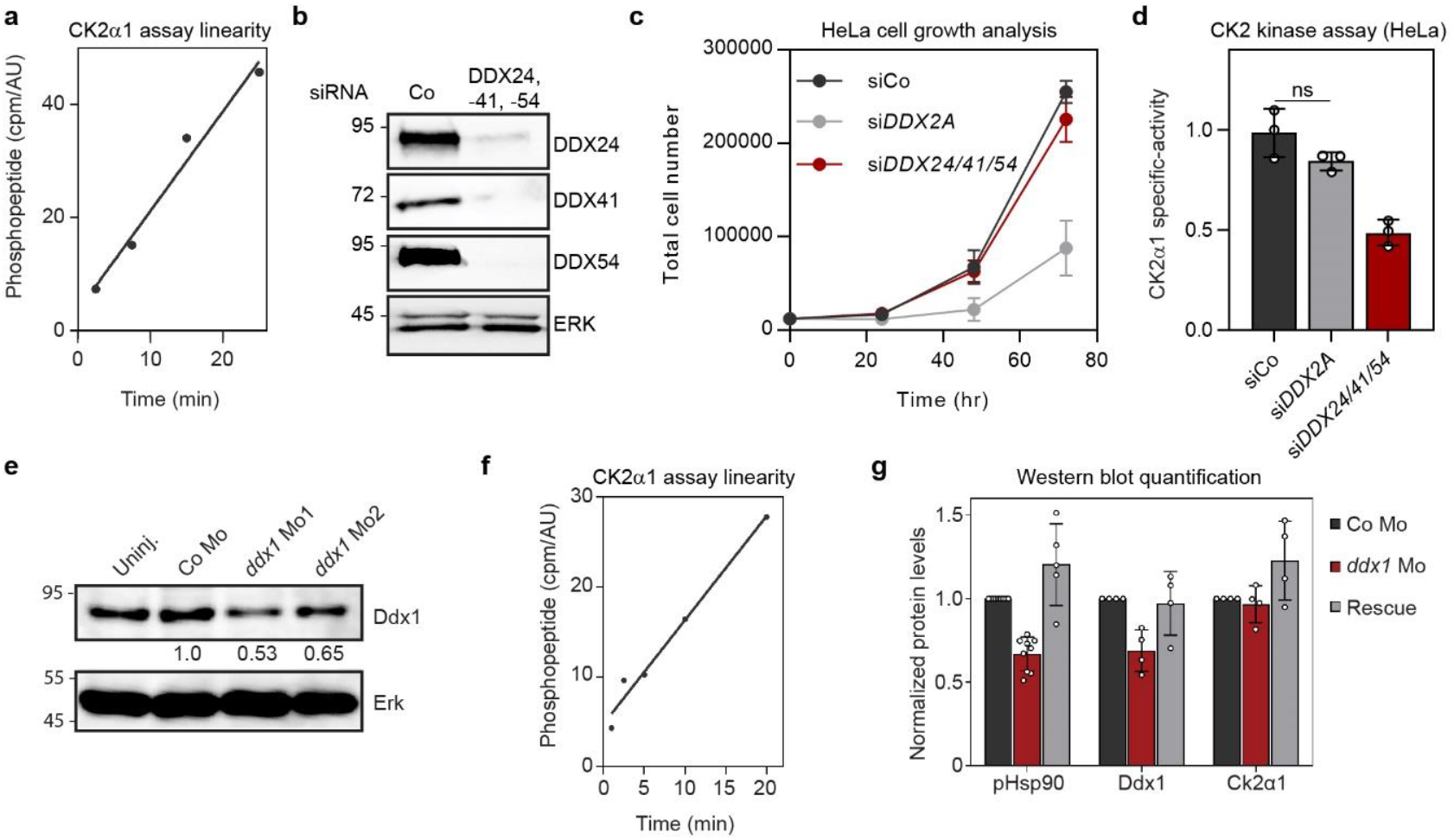
DDX proteins bind to and regulate CK2α1/α2 activity. **a**, Related to Main Fig. 2c. *In vitro* kinase assay with CK2α1 immunopurified from HeLa cells. Phosphopeptide (cpm) was normalized to the kinase amount used (AU). n = 1. **b**, Related to Main Fig. 2c. Western blot of HeLa cells treated with the siRNA confirming knockdown of the corresponding DDX. **c**, Cell growth analysis of HeLa cells after transfection with the indicated siRNA. n = 3 biological replicates. **d**, CK2α1 specific activity as determined from immunopurified and normalized CK2α1 protein from HeLa cells after siRNA knockdown of the indicated siRNA. n = 3 biological replicates. **e**, Western blot of Ddx1 in *X. tropicalis* embryos (stage 18) after injection of 20 ng of Mo1 or 40 ng Mo2. Knockdown efficiency is indicated. **f**, Related to Main Fig. 2i-j. *In vitro* kinase assay with Ck2α1 immunopurified from *X. tropicalis* embryos (stage 18). Phosphopeptide (cpm) was normalized to the kinase amount used (AU). n = 1. **g**, Related to Main Fig. 2k. Quantification of pHsp90, Ddx1, and Ck2α1 protein levels in *X. tropicalis* embryos (stage 18) after ddx1 Mo injection. n ≥ 4 biological replicates. Data are means ± SD with one-way ANOVA (**d, g**).

**Extended Data Fig.3.**
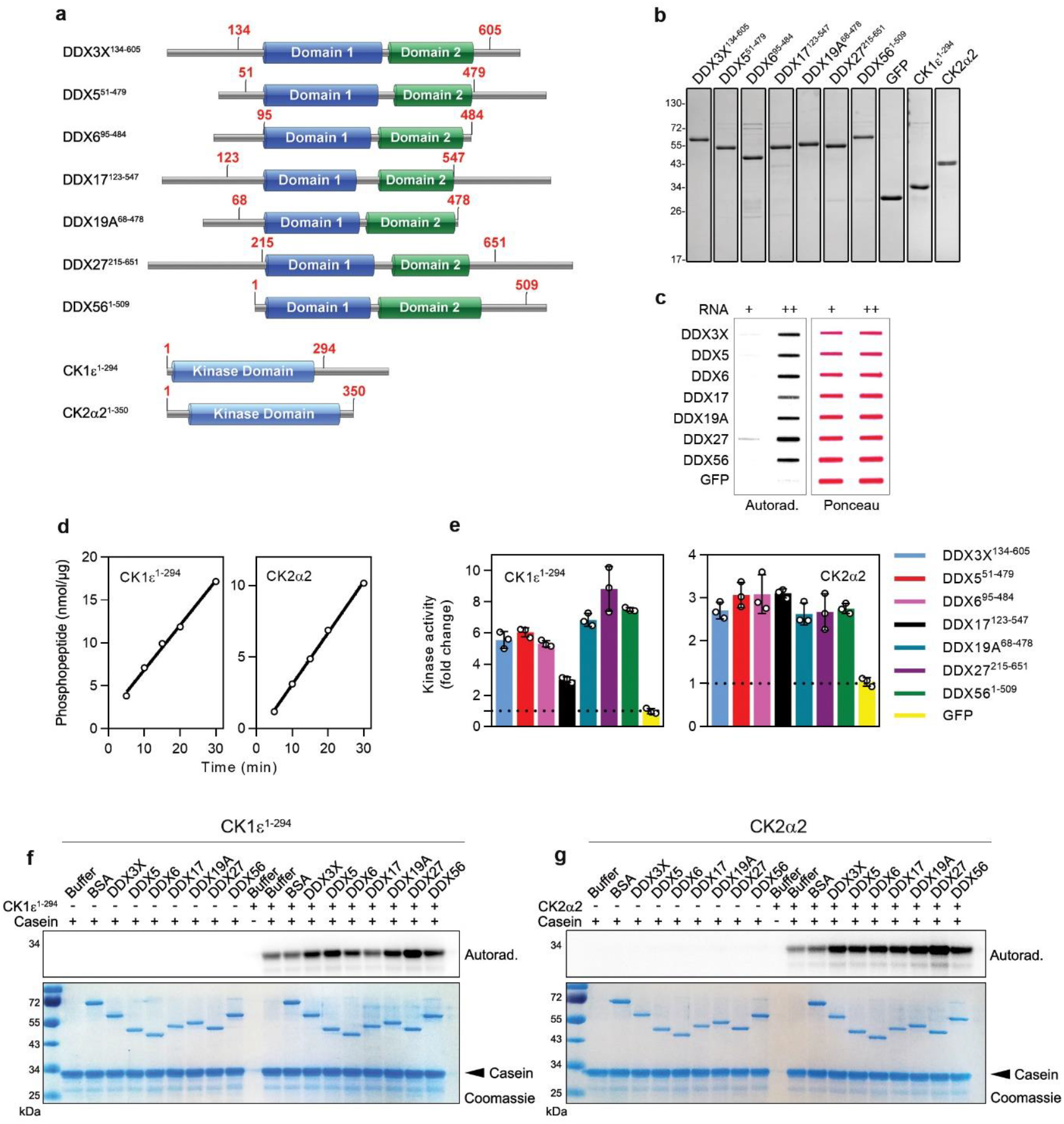
Expression, purification and functional characterization of DDX core domain proteins and protein kinases. **a**, Cartoon of DDX core domain and protein kinase constructs used in this study created using IBS 1.0.3^32^. Red amino acid numbers indicate the boundaries of each DDX core domain and protein kinase construct. **b**, Coomassie stained SDS-PAGE gel of recombinant DDX core domains and protein kinases expressed and purified from *E. coli*. **c**, RNA binding assay with DDX proteins. Autoradiogram of DDX core domain- or GFP-loaded slot blots incubated with ^32^P-polyA^-^-RNA; + = 0.1 ng/μl, ++ = 1 ng/μl. **d**, *In vitro kinase* assays with the indicated protein kinase and peptide substrate. [Peptide substrate] = 100 µM for both CK1ε ^1-294^ and CK2α2. n = 1. **e**, *In vitro* kinase assays for CK1ε ^1-294^ and CK2α2 and the indicated DDX core domain proteins or GFP. [Peptide substrate] = 200 µM for CK2α2. Kinase activity is normalized to buffer only. Data are means ± SD; n = 3. **f-g**, Stimulation of casein phosphorylation by CK1ε ^1-294^ (**f**) or CK2α2 (**g**) by DDX core domain proteins. After *in vitro* kinase assays with 32P-γATP, proteins were analysed by SDS-PAGE followed by Coomassie stain (bottom) and autoradiography (top).

**Extended Data Fig.4.**
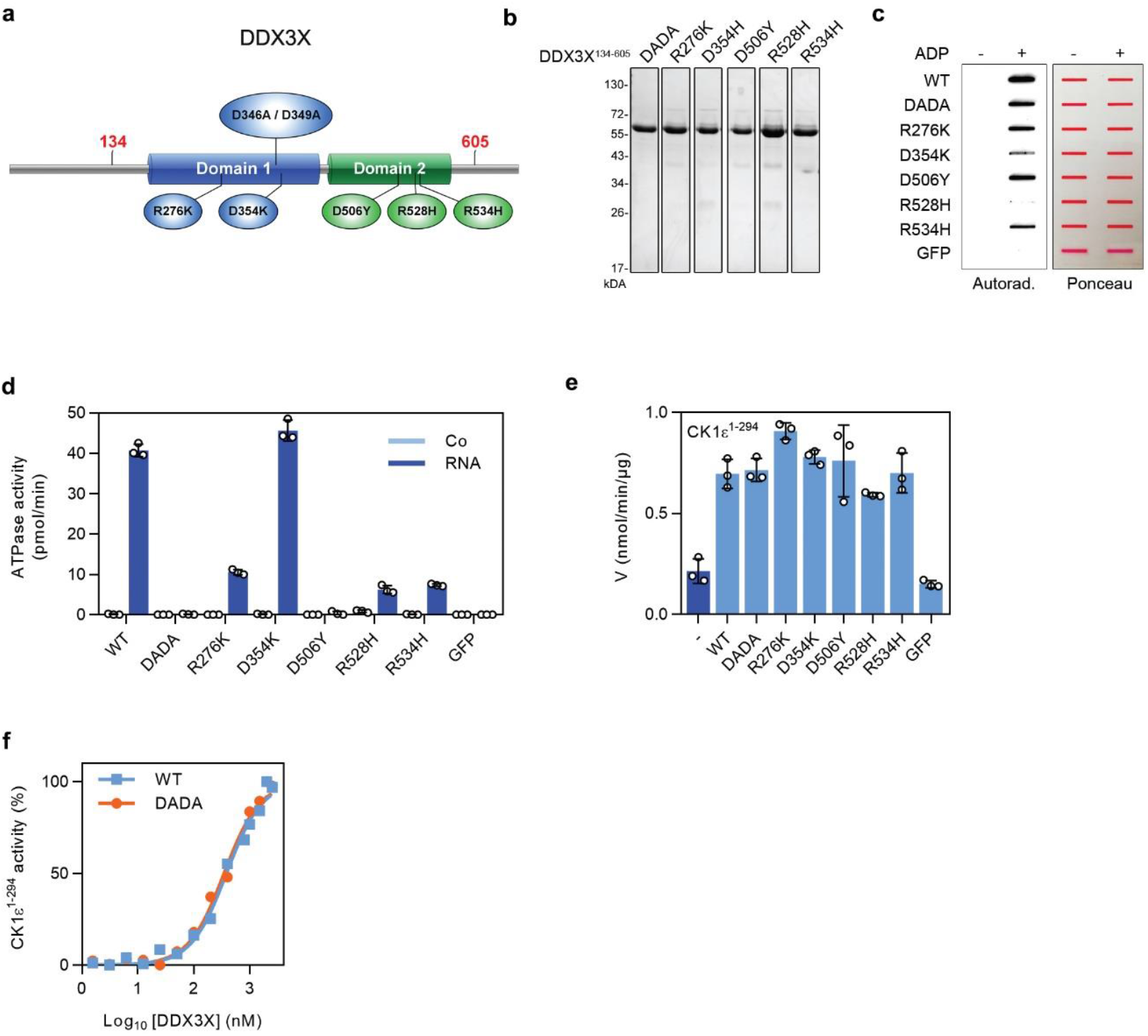
DDX helicase activity is not required for CK1ε ^1-294^ kinase activity stimulation. **a**, Cartoon of DDX3X highlighting amino acids of mutants used for *in vitro* kinase assays created using IBS 1.0.3^32^. ATPase dead form D346A/D349A (DDX3X^DADA^) mutant and point mutants designed after mutations found in Wnt-driven medulloblastoma patients are indicated. Red amino acid numbers indicate the boundaries of the DDX3X core domain construct. **b**, Coomassie stained SDS-PAGE gel of purified recombinant DDX3X mutant proteins expressed in *E. coli*. **c**, ADP binding assay with DDX3X mutants. Autoradiogram of DDX3X mutants- or GFP-loaded slot blots incubated with ^32^P-αADP. **d**, ATPase assay with DDX3X mutants with or without polyA^-^-RNA. Data are means ± SD; n = 3. **e**, *In vitro* kinase assay with recombinant CK1ε ^1-294^ and the indicated DDX3X mutants. Data are means ± SD; n = 3. **f**, Dose response curve of CK1ε ^1-294^ kinase activity with increasing concentrations of DDX3X^134- 605^ (WT) and DDX3X D346A/D349A (DADA).

**Extended Data Fig.5.**
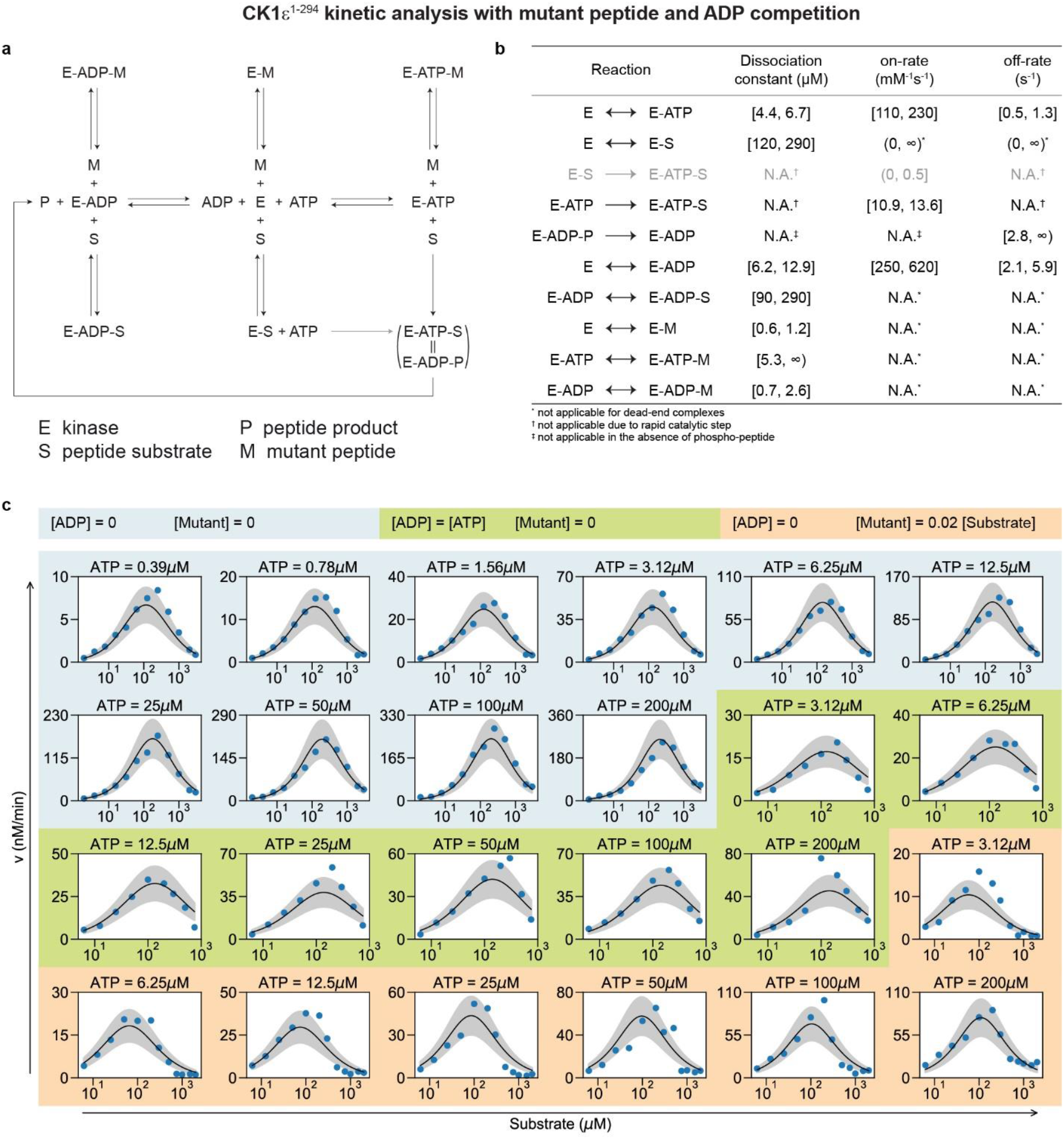
Fit of the enzyme-kinetic model for CK1ε ^1-294^. Related to Fig. 4. **a**, Reaction scheme of the best-fitting model, including competitive inhibition by non-phosphorylatable mutant peptide. **b**, Best-fit parameters, with 95% confidence bounds (∞, no upper confidence bound was found, i.e., the model fit remains good with the respective parameter becoming very large). **c**, Simultaneous fitting of the model to the reaction velocities measured for a concentration matrix of peptide substrate, ATP, ADP and non-phosphorylatable mutant peptide, as indicated. Blue dots, experimental data (n = 1); black curves, best model fit; grey shades, ±1 SD error bands.

**Extended Data Fig.6.**
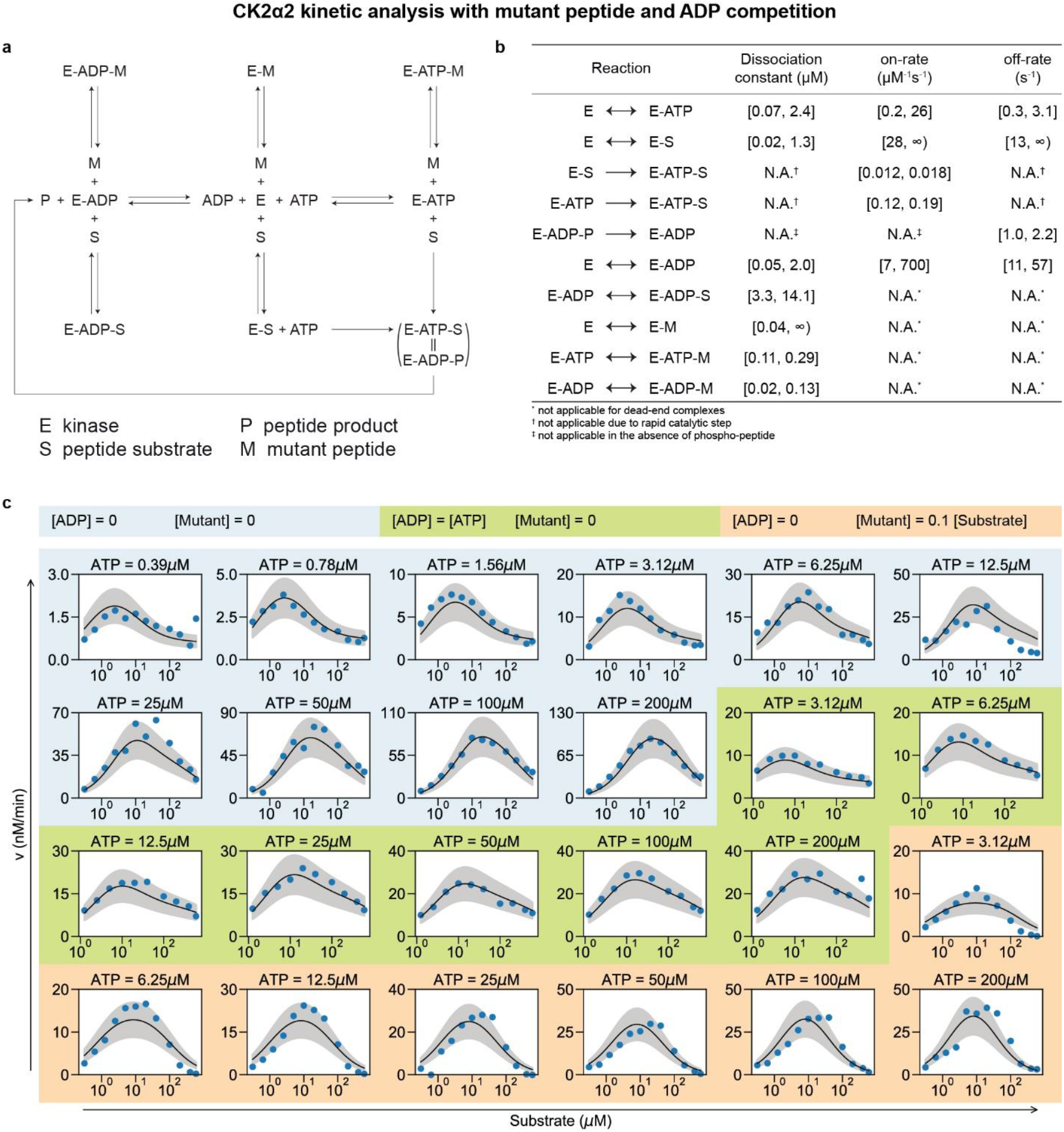
Fit of the enzyme-kinetic model for CK2α2. Related to Fig. 4. **a**, Reaction scheme of the best-fitting model, including competitive inhibition by non-phosphorylatable mutant peptide. **b**, Best-fit parameters, with 95% confidence bounds [0 (∞), no lower (upper) confidence bound was found, i.e., the model fit remains good with the respective parameter becoming very small (large)]. **c**, Simultaneous fitting of the model to the reaction velocities measured for a concentration matrix of peptide substrate, ATP, ADP and non-phosphorylatable mutant peptide, as indicated. Blue dots, experimental data (n = 1); black curves, best model fit; grey shades, ±1 SD error bands.

**Extended Data Fig.7.**
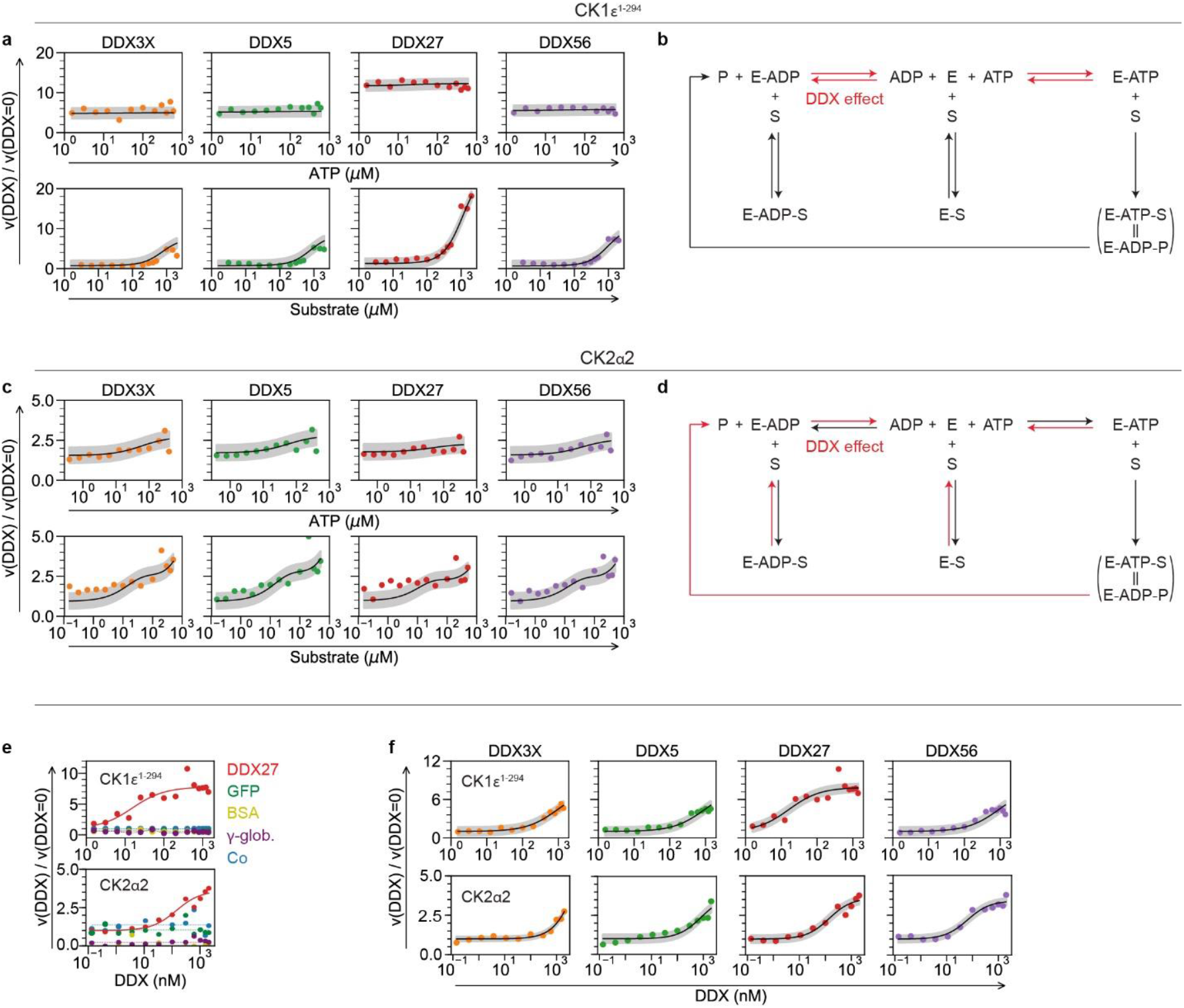
Model selection to detect activation mechanism by DDX proteins. **a**, Fit of the best model for CK1ε ^1-294^ to measured reaction velocity data in the presence of DDX core domain proteins, as indicated. **b**, Scheme of the selected best model for CK1ε ^1-294^ (Fig. 4a, g). **c**, Fit of best model for CK2α2 to measured reaction velocity data in the presence of DDX proteins, as indicated. **d**, Scheme of the selected best model for CK2α2 (Fig. 4a, g). **e**, Addition of DDX27^215-651^ but not of control proteins (GFP, BSA, γ-globulin), increases the reaction velocities of CK1ε ^1-294^ and CK2α2 in a dose-dependent manner. **f**, The dose effects of DDX proteins on reaction velocities of CK1ε ^1-294^ and CK2α2 are reproduced by the corresponding models. Experimental data show one representative experiment of n = 2 (**a, c, e, f**).

**EExtended Data Fig. 8.**
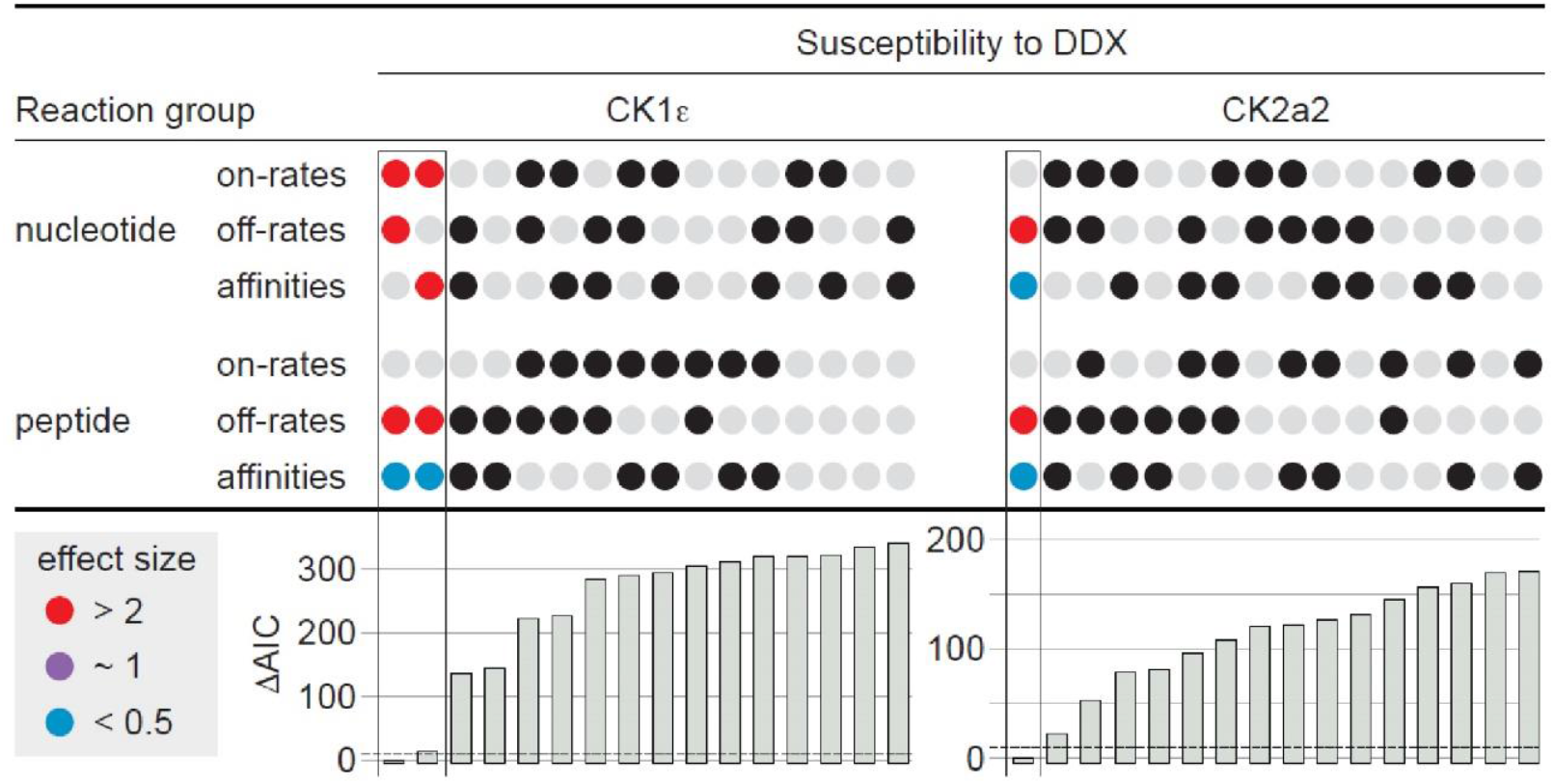
Allosteric model for DDX mediated kinase stimulation predicts accelerated ADP release. Prediction of the mechanism of DDX action by statistical model selection considering allosteric substrate binding rather than by forming a dead-end complex as in (Fig. 4g). DDX proteins were allowed to affect on- or off-rates for adenine nucleotides (ATP/ADP) or peptide, either as single parameters or in any combination, constrained by equal effect sizes for a given pair of on- and off-rates when both were allowed to change upon DDX binding (upper panels). The best fits of each model to the measured velocity data were ranked by the Akaike information criterion (AIC, lower panel). The effect sizes for DDX27^215-651^ are displayed for the models close to or passing the standard selection threshold (ΔAIC < 10). Note that the best fitting models also support accelerated ADP release by DDX27^215-651^.

## Notes

### Competing Interest Statement

The authors have declared no competing interest.

## References

1. Henn, A., Bradley, M.J. & De La Cruz, E.M. ATP utilization and RNA conformational rearrangement by DEAD-box proteins. Annu Rev Biophys 41, 247–67 (2012).

2. Linder, P. & Jankowsky, E. From unwinding to clamping - the DEAD box RNA helicase family. Nat Rev Mol Cell Biol 12, 505–16 (2011).

3. Cruciat, C.M. et al. RNA helicase DDX3 is a regulatory subunit of casein kinase 1 in Wnt-β-catenin signaling. Science 339, 1436–41 (2013).

4. Dolde, C. et al. A CK1 FRET biosensor reveals that DDX3X is an essential activator of CK1ε. J Cell Sci 131(2018).

5. Bol, G.M. et al. Targeting DDX3 with a small molecule inhibitor for lung cancer therapy. EMBO Mol Med 7, 648–69 (2015).

6. Heerma van Voss, M.R. et al. Identification of the DEAD box RNA helicase DDX3 as a therapeutic target in colorectal cancer. Oncotarget 6, 28312–26 (2015).

7. Snijders Blok, L. et al. Mutations in DDX3X Are a Common Cause of Unexplained Intellectual Disability with Gender-Specific Effects on Wnt Signaling. Am J Hum Genet 97, 343–52 (2015).

8. Lauinger, L., Diernfellner, A., Falk, S. & Brunner, M. The RNA helicase FRH is an ATP-dependent regulator of CK1a in the circadian clock of Neurospora crassa. Nat Commun 5, 3598 (2014).

9. Rogers, G.W., Komar, A.A. & Merrick, W.C. eIF4A: the godfather of the DEAD box helicases. Prog Nucleic Acid Res Mol Biol 72, 307–31 (2002).

10. Lou, D.Y. et al. The alpha catalytic subunit of protein kinase CK2 is required for mouse embryonic development. Mol Cell Biol 28, 131–9 (2008).

11. Rusin, S.F., Adamo, M.E. & Kettenbach, A.N. Identification of Candidate Casein Kinase 2 Substrates in Mitosis by Quantitative Phosphoproteomics. Front Cell Dev Biol 5, 97 (2017).

12. Lees-Miller, S.P. & Anderson, C.W. Two human 90-kDa heat shock proteins are phosphorylated in vivo at conserved serines that are phosphorylated in vitro by casein kinase II. J Biol Chem 264, 2431–7 (1989).

13. Hildebrandt, M.R., Germain, D.R., Monckton, E.A., Brun, M. & Godbout, R. Ddx1 knockout results in transgenerational wild-type lethality in mice. Sci Rep 5, 9829 (2015).

14. Pugh, T.J. et al. Medulloblastoma exome sequencing uncovers subtype-specific somatic mutations. Nature 488, 106–10 (2012).

15. Garbelli, A., Beermann, S., Di Cicco, G., Dietrich, U. & Maga, G. A motif unique to the human DEAD-box protein DDX3 is important for nucleic acid binding, ATP hydrolysis, RNA/DNA unwinding and HIV-1 replication. PLoS One 6, e19810 (2011).

16. Floor, S.N., Condon, K.J., Sharma, D., Jankowsky, E. & Doudna, J.A. Autoinhibitory Interdomain Interactions and Subfamily-specific Extensions Redefine the Catalytic Core of the Human DEAD-box Protein DDX3. J Biol Chem 291, 2412–21 (2016).

17. Song, H. & Ji, X. The mechanism of RNA duplex recognition and unwinding by DEAD-box helicase DDX3X. Nat Commun 10, 3085 (2019).

18. Cornish-Bowden, A. Fundamentals of Enzyme Kinetics. (Portland Press, London, 1995).

19. Adams, J.A. Kinetic and catalytic mechanisms of protein kinases. Chem Rev 101, 2271–90 (2001).

20. Akaike, H. Information Theory and an Extension of the Maximum Likelihood Principle. 267–281 (Springer, Springer Series in Statistics, 1973).

21. Burnham, K. & Anderson, D. Model selection and multimodel inference: a practical information-theoretic approach. xxvi, 488 p. (Springer, New York, ed. 2nd, 2002, 2002).

22. Ni, Q., Shaffer, J. & Adams, J.A. Insights into nucleotide binding in protein kinase A using fluorescent adenosine derivatives. Protein Sci 9, 1818–27 (2000).

23. Aubol, B.E., Plocinik, R.M., McGlone, M.L. & Adams, J.A. Nucleotide release sequences in the protein kinase SRPK1 accelerate substrate phosphorylation. Biochemistry 51, 6584–94 (2012).

24. Bracher, A. & Verghese, J. The nucleotide exchange factors of Hsp70 molecular chaperones. Front Mol Biosci 2, 10 (2015).

25. Andréasson, C., Fiaux, J., Rampelt, H., Druffel-Augustin, S. & Bukau, B. Insights into the structural dynamics of the Hsp110-Hsp70 interaction reveal the mechanism for nucleotide exchange activity. Proc Natl Acad Sci U S A 105, 16519–24 (2008).

26. Endicott, J.A., Noble, M.E. & Johnson, L.N. The structural basis for control of eukaryotic protein kinases. Annu Rev Biochem 81, 587–613 (2012).

27. Zhou, J. & Adams, J.A. Participation of ADP dissociation in the rate-determining step in cAMP-dependent protein kinase. Biochemistry 36, 15733–8 (1997).

28. Burke, K.A., Janke, A.M., Rhine, C.L. & Fawzi, N.L. Residue-by-Residue View of In Vitro FUS Granules that Bind the C-Terminal Domain of RNA Polymerase II. Mol Cell 60, 231–41 (2015).

29. Rai, A.K., Chen, J.X., Selbach, M. & Pelkmans, L. Kinase-controlled phase transition of membraneless organelles in mitosis. Nature 559, 211–216 (2018).

30. Hondele, M. et al. DEAD-box ATPases are global regulators of phase-separated organelles. Nature 573, 144–148 (2019).

31. Cai, W. et al. Wanted DEAD/H or Alive: Helicases Winding Up in Cancers. J Natl Cancer Inst 109(2017).

32. Liu, W. et al. IBS: an illustrator for the presentation and visualization of biological sequences. Bioinformatics 31, 3359–61 (2015).

