## Supplementary Information for "DEAD-box RNA Helicases Act as Nucleotide Exchange Factors for Casein Kinase 2"

### Supplementary Materials and Methods

### Constructs

Human *DDX1* (EL735960), *DDX5* (EL733383), *DDX6* (BC065007), *DDX17* (BC000595), *DDX19A* (BC), *DDX20* (BC011556), *DDX27* (EL733496), *DDX39A* (NM_005804), *DDX-41* (BC015476), *DDX50* (BC000272), *DDX56* (BC001235), *CSNK2A1* (BC011668), and *CSNK2A2* (BC008812) cDNAs were obtained from the German Cancer Research Center (DKFZ) clone collection. Human *DDX3X* (BC 063374) and *CSNK1E* (BC006490) cDNA was obtained from imaGenes. Mouse *CSNK1A* (BC048081), human *CSNK1D* (BC003558) and *NME2* (BC095458) cDNAs were obtained from Source BioScience. DDX3X medulloblastoma associated mutations were a kind gift from Beat Suter[4]. The DDX3X D346A/D349A (DDX3X$^{\mathrm{DADA}}$) construct was a kind gift from Giovanni Maga[15]. For mammalian expression, tagged constructs were generated by inserting the respective full-length cDNA into pCS2+ vector containing an N- or C-terminal V5, Myc or FLAG tag. For bacterial expression, DDX, CK1$\varepsilon^{1\text{-}294}$ and CK2$\alpha$2 protein kinases were cloned into modified pET-based vector with N-terminus 6xHis-tag[33]. DDX3X$^{134\text{-}605}$ core domain was generated as described[16]. Medulloblastoma associated mutants, DADA mutations, and Alanine scanning mutations were introduced in the DDX3X$^{132\text{-}605}$ background. DDX5$^{51\text{-}479}$, DDX6$^{95\text{-}484}$, DDX17$^{123\text{-}547}$, DDX19A$^{68\text{-}478}$, DDX27$^{215\text{-}651}$, and DDX56$^{1\text{-}509}$ core domains were generated by amplifying the appropriate sequences predicted to be globular by the Crystallization Construct Designer web tool (https://ccd.rhpc.nki.nl/) and cloned into modified pET-based vector with N-terminus 6xHis-tag[33]. *Xenopus ck2*$\alpha$, and *ck2*$\beta$ were a kind gift from Isabel Dominguez[34]. *Xenopus ddx1* was cloned from *X. tropicalis* (stage 40) cDNA into pCS2+ vector.

### Xenopus experiments

All *Xenopus* experiments were approved by the state of Baden-Württemberg (Germany) and performed according to federal guidelines. *X. laevis* (J-strain, inbred) were obtained from Nasco (USA) and EXRC (England). *X. tropicalis* (Xtr.NigerianSt549, inbred strain) were obtained from Nasco (USA), EXRC (England), NXR (USA) and CRB (France). Frogs were kept in a *Xenopus* facility designed by Tecniplast with regulated water parameters, temperature (18 °C for *X. laevis* / 25 °C for *X. tropicalis*), and day-night cycles (12/12h). Egg laying was induced by injection of 600 U of hCG (Aska Pharmaceuticals) in *X. laevis* and 120 U in *X. tropicalis*. *In vitro* fertilization and embryo injections were carried out as described[35].

### Morpholino knockdowns

Two antisense Morpholino oligonucleotides (Mo) targeting the *X. tropicalis ddx1* gene (ENSXETG00000016652.1) were designed for specific Ddx1 loss-of-function experiments. *ddx1* Mo1: 5' CCA TGA ACA ATC CCG CTG TAC CTG A 3', *ddx1* Mo2: 5' CAG CTG ACT CGG AAG TAG TAA CCG A 3'. For gain-of-function studies, sense mRNA was generated using the SP6 Amplicon MegaScript Kit (Ambion). *X. tropicalis* embryos were injected at stage 2 - 4 with 20 ng *ddx1* Mo (splice site Mo) and/or 100 pg *ddx1* mRNA unless stated otherwise. Control embryos were injected with an equal dose of standard control Mo (GeneTools) and/or PPL mRNA.

### Expression and purification of recombinant proteins from E. coli

$CK1\varepsilon^{1\text{-}294}$, CK2α2, DDX core domains and DDX3X mutants were expressed in *E. coli* Rosetta$^{TM}$(DE3)pLysS (Novagen) with auto-induction medium system. Cells were grown at 37 °C in 500 ml ZY complete medium[36] to an $OD_{600}$ of 0.8, then the temperature was lowered to 20 °C and growth continued for 18 h at 220 rpm. Cells were harvested, washed in cold PBS and cell pellet flash frozen in liquid nitrogen and stored at -20 °C until use. Cell pellet was dissolved in 10 ml/g of Lysis Buffer A (20 mM HEPES pH 8.0, 500 mM NaCl, 5 mM $MgCl_2$, 0.2 % NP-40, 10 mM Imidazole, 5 mM β-mercaptoethanol, 10 U/ml Benzonase®, 0.5 mg/ml Lysozyme) supplemented with c0mplete protease inhibitor cocktail (Roche), and mechanically disrupted by Microfluidizer® (M1-10L, Microfluidics Inc., USA). The lysate was centrifuged at 23,000 rpm (JA25.50 fixed angle rotor, Beckman). The supernatant was filtered through a 0.45 µm PVDF membrane filter (Millipore) and incubated with Ni-NTA beads (Quiagen). Beads were washed with 5 column volumes of Wash Buffer A (20 mM HEPES pH 8.0, 500 mM NaCl, 5 mM $MgCl_2$, 0.2 % NP-40, 10 mM Imidazole, 2 mM β-mercaptoethanol), Wash Buffer B (20 mM HEPES pH 8.0, 1 M NaCl, 5 mM $MgCl_2$, 10 mM Imidazole, 2 mM β-mercaptoethanol), Wash Buffer C (20 mM HEPES pH 8.0, 500 mM NaCl, 5 mM $MgCl_2$, 20 mM Imidazole, 2 mM β-mercaptoethanol), and eluted in 2.5 column volumes of Elution Buffer A (20 mM HEPES pH 8.0, 500 mM NaCl, 5 mM $MgCl_2$, 330 mM Imidazole, 2 mM β-mercaptoethanol). The elution buffer was exchanged to Storage Buffer (20 mM HEPES pH 8.0, 500 mM NaCl, 5 mM $MgCl_2$, 1 mM DTT, 10 % Glycerol) by passing the proteins through a PD-10 column (GE Healthcare), concentrated with centrifugal filter units (Millipore) and purity was assessed by SDS-PAGE and Coomassie stain

(Quick Coomassie® stain, SERVA). Large scale purifications were further purified by size exclusion chromatography on a Superdex 200 26/60 column (GE Healthcare) using an ÄKTAprime plus system (GE Healthcare). 50 µM protein aliquots were flash frozen and stored at -80 °C.

### Expression and purification of recombinant DDX1, CK1, CK2 and NME2 from HEK293T cells

Recombinant FLAG-DDX1 and FLAG-tagged protein kinases were expressed and purified from HEK293T cells. 15 cm dishes were transfected with 20 µg of FLAG-tagged protein vector and expression continued for 48 h. Cells were harvested in cold PBS and pellet lysed in 10 ml/g of Lysis Buffer B (50 mM HEPES, pH 8.0, 300 mM NaCl, 5 mM $MgCl_2$, 0.8 % Triton X-100, 10 mM NaF, 0.2 % Sodium deoxycholate, 0.05 % SDS, 5 mM β-mercaptoethanol, 1x c0mplete protease inhibitor cocktail (Roche)). Cleared lysate was incubated with FLAG-M2 affinity gel (SIGMA) for 16 h at 4 °C in constant rotation. Beads were washed once with Wash Buffer D (30 mM HEPES pH 8.0, 300 mM NaCl, 5 mM $MgCl_2$, 0.2 % NP-40, 2 mM β-Mercaptoethanol), twice with Wash Buffer E (20 mM HEPES pH 8.0, 500 mM NaCl, 5 mM $MgCl_2$, 2 mM β-mercaptoethanol), and twice with Elution Buffer B (30 mM HEPES pH 8.0, 300 mM NaCl, 0.1 % Tween-20, 1 mM $MgCl_2$, 20 % Glycerol, 2 mM β-mercaptoethanol). Proteins were eluted from FLAG-M2 beads by FLAG peptide competition using 200 µg/ml FLAG peptide (SIGMA) in Elution Buffer B. Purity of eluted proteins was assessed by SDS-PAGE followed by a Coomassie stain, quantification and storage in aliquots at -80 °C until use.

### In vitro binding assay

For *in vitro* binding assays, Myc-tagged DDX proteins were expressed and immunopurified from HEK293T cells. 15 cm dishes were transfected with 20 µg of Myc-tagged DDX protein vector and expression continued for 48 h. Cells were harvested in cold PBS and lysed in 10 ml/g of Lysis Buffer B. Cleared lysate was incubated with mouse anti-Myc (9E10) antibody followed by incubation with protein G agarose beads (Santa Cruz) for 16 h at 4 °C with constant rotation. Beads were washed once with Wash Buffer D, twice withWash Buffer E, and once with Binding Buffer (30 mM HEPES pH 8.0, 100 mM NaCl, 0.1 % Tween-20, 1 mM $MgCl_2$, 2 mM β-Mercaptoethanol). Myc-DDX conjugated beads were blocked in 1 % BSA in Binding Buffer for 1 h at 4 °C and washed in Binding Buffer supplemented with 0.1 % BSA. Finally, the beads were incubated with 400 ng of FLAG-tagged protein kinases purified as described above in 0.1 % BSA in Binding Buffer.

After 3 h incubation at 4 °C beads were washed 4 times in Wash Buffer F (30 mM HEPES pH 8.0, 100 mM NaCl, 0.1% TritonX-100, 1 mM $MgCl_2$, 2 mM β-mercaptoethanol) and binding analyzed by SDS-PAGE followed by western blot analysis.

### Co-immunoprecipitation (CoIP)

For CoIP between CK2α1 and DDX24, -41, and -54, HeLa cells were seeded in a 10 cm dish and harvested after reaching full confluence by trypsinisation. Cells were washed once in PBS and lysed in 2 ml of IP-Lysis Buffer (50 mM TRIS-HCl pH 8.0, 150 mM NaCl, 0.1 % Triton X-100, 1x c0mplete protease inhibitor cocktail (Roche)). The lysate was split in 4 x 500 µl aliquots. CK2α1 was immunopurified by adding 2 µg of α-CK2α1 (Abfrontier, LF-MA0223) and 10 µl of Protein A magnetic beads (NEB) to two of the aliquots. Control IPs were performed by using 2 µg of IgG raised in the same species. After incubation at 4 °C overnight, beads were washed 3x in IP-Lysis buffer, and boiled in either 25 µl or 50 µl of Laemmli buffer for 5 minutes at 99 °C. After Western blotting, proteins were detected using the following antibodies: CK2α1 (Bethyl, A300-198A), CK2β (Bethyl, A301-984A), DDX24 (Bethyl, A300-698A), DDX41 (Bethyl, A301-050A), DDX54 (Bethyl, A3001-313A), DDX56 (Abcam, ab115158) all at a dilution of 1:1000. For CoIP between Ddx1 and Ck2α, 800 *X. tropicalis* embryos were collected at stage 18 and after removal of excess liquid lysed in one volume of XT-Lysis Buffer (50 mM TRIS-HCl pH 8.0, 145 mM NaCl, 0.05 % NP-40, 5 % Glycerol, 1x c0mplete protease inhibitor cocktail (Roche), 1x Phosphatase Inhibitor (Thermo Fisher)). Lipids were removed by two times Freon (CFC-113, Honeywell 34874) extraction, and the lysate split in four equal parts. Ddx1 was immuno-purified using 1:1000 α-DDX1 (Bethly, A300-521A), and Ck2α using 1:1000 α-CK2α (Abfrontier, LF-MA0223) and 100 µl of Protein A agarose beads (Santa Cruz) overnight at 4 °C. Control IPs were performed with equal amounts of IgG raised in the same species. Beads were washed 3 x in XT-Lysis buffer, and boiled in Laemmli Buffer for 5 minutes at 99 °C. After Western blotting, proteins were detected using the same antibodies.

### Wnt reporter assay (TopFlash)

HEK293T cells were seeded at 50,000 cells per well to a 96-well plate and transfected with 50 nM siDDX3. The following day, each well was transfected with 15 ng of Super-TopFlash, 3 ng of Renilla, and the indicated DDX constructs. Plasmid amounts were adjusted to achieve comparable expression levels, ranging from 30 to 50 ng per well. 36 hours post DNA transfection, Wnt3a conditioned media was added to each well and incubated overnight. The cells were lysed in 1x passive lysis buffer (Promega) for 10 minutes at 4 °C. Luciferase and Renilla activities were monitored, using the dual Luciferase Assay Kit (Promega) and SPARK plate reader (Tecan).

### Kinase Assay, Kinase Immunoprecipitation, and ELISA

Kinase assays were performed by filter binding with $^{32}$P-γATP (PerkinElmer NEG-502A001MC, 3000 Ci/mmol) and specific peptide substrate (CK1tide: RRKDLHDDE-EDEAMSITA CK2tide: RRRADDSDDDDD). A non-phosphorylatable peptide substrate mutant form was synthesized by replacing the Serine (highlighted in red) by Alanine. Preliminary experiments were performed to determine optimal assay conditions for each protein kinase. All kinase reactions were carried out in the linear phase of the reaction progress curve (initial rate). For Figure S1, human recombinant CK1 full-length isoforms were obtained from Invitrogen. Human recombinant full-length DDX6, DDX20, DDX39A, DDX41, DDX50 and DDX56 were obtained from Abnova. CK1 kinase assays were performed in 30 µl in 1.5 ml tubes at 30 °C for 15 minutes in Kinase Buffer A (30 mM HEPES-KOH pH 7.7, 10 mM $MgCl_2$, 0.02 µg/ml BSA, 1 mM DTT). For CK2α2 Kinase Buffer B (30 mM TRIS-HCl pH 8.5, 10 mM $MgCl_2$, 0.02 µg/ml BSA, 1 mM DTT) was used. Control proteins or DDX proteins were incubated with CK1 or CK2 and their corresponding specific peptide substrate. If not indicated otherwise, 2.5 nM kinase, 500 nM DDX or control protein, and 1 mM (CK1) or 0.1 mM (CK2α2) peptide substrate were used. Assay components were premixed and reactions were started by the addition of 50 - 100 µM ATP containing 1 µCi of radiolabeled $^{32}$P-γATP and stopped by pipetting 10 µl onto p81 phosphocellulose filter paper (ST Vincent's Institute). In Figure 4L, poly-uridine (Sigma #P9528) was added to the kinase/DDX mix to allow for pre-incubation before the reaction was started. P81 filter paper was washed 4 times in 0.5 % ortho-phosphoric acid, once in 100 % ethanol, dried and radioactivity was detected by Cherenkov radiation in a scintillation counter (Packard Tri-Carb 2100 TR). Kinase specific activity was expressed as nmol of phosphate transferred from ATP to peptide substrate per minute per µg of kinase (nmol/min/µg). Casein phosphorylation assays were performed as described above in presence of 500 nM DDX proteins or BSA, with 2.5 nM CK1ε and CK2α2 for 15 min with 0.1 mg/ml dephosphorylated α-casein (Sigma, C8032) instead of peptide substrate. After the kinase reaction, the sample was analyzed by 12.5 % SDS-PAGE followed by staining with Quick Coomassie® (Serva) and phosphorimager (Sapphire™ Biomolecular Imager (Azure Biosystems)) of the dried gel.

For kinase assay from *X. tropicalis*, embryos were injected with either 20 ng control or *ddx1* Morpholino. At stage 18, 20 embryos per replicate were collected and lysed in XT-Lysis

Buffer. Lipids were removed by Freon extraction (CFC-113, Honeywell 34874). The lysate was subjected to Ck2α pulldown using 0.5 µg α-CK2α (Abfrontier, LF-MA0223) and 7 µl Protein A magnetic beads (NEB) overnight at 4 °C. Beads were washed 3x in XT-Lysis Buffer and 2x in XT-Kinase Buffer (30 mM HEPES-KOH pH 7.7, 5 mM $MgCl_2$, 1 mM DTT, 0.1 % BSA). Beads were resuspended in 30 µl XT-Kinase Buffer supplemented with 0.5 mM CK2tide, 100 µM ATP with 1 µCi radiolabeled $^{32}$P-γATP and the kinase reaction was carried out for 20 minutes at 37 °C while shaking (1100 rpm). 10 µl of the kinase reaction was spotted onto P81 filter paper and processed as described above. To quantitate the kinase bound to the beads, the kinase was eluted in 50 µl of 200 mM Glycine Buffer pH 2.6. After neutralization with an equal volume of 1 M TRIS-HCl pH 8.0, the eluate was supplemented with 100 µl of 2x Coating Buffer (100 mM $NaHCO_3$, pH 9.6) and allowed to bind to an ELISA plate (Greiner) overnight at 4 °C. After washing 2x and blocking for 30 minutes in 4 % BSA, the wells were incubated with α-CK2α1 (Bethyl, A300-198A, 1:1000 in 4 % BSA + TBS-T) and after washing 5x with TBST-T, decorated with α-rabbit-HRP (Dianova, 1:1000 in 4 % BSA + TBS-T). After 5x washing with TBS-T, HRP activity was detected using the fluorometric detection kit QuantaBluTM (Thermo Fisher).

For kinase assays from mammalian cell lines, HEK293T or HeLa cells were plated in 24-well plates (50.000 cells per well) and transfected with 50 nM of the corresponding siRNA. Three days post transfection, cells were harvested in 500 µl of HEK-Lysis Buffer (50 mM HEPES-KOH pH 7.7, 150 mM NaCl, 1 % NP-40, 5 mM $MgCl_2$, 1x c0mplete protease inhibitor cocktail (Roche), 1x Phosphatase Inhibitor (Thermo Fisher)). For the CK1ε kinase assay from HEK293T cells, the lysate of one well was used per sample, for the CK2α1 kinase assay from HeLa cells, 25 % of cell lysate per 24-well were used for the kinase assay. Peptide concentrations for CK1tide were 1 mM, for CK2tide 0.5 mM if not indicated otherwise. Kinase assay and ELISA quantification was performed as described above for *X. tropicalis*. For determination of $K_m$[S], $K_m$[ATP], $K_i$, and $V_{max}$, the reaction velocity data were fitted using non-linear regression to the Michaelis-Menten equation or to the substrate inhibition equation (equation 5.43)[37]

$$v = V_{max} * [S]/(K_m + [S] * (1 + [S]/K_i)) \tag{1a}$$

where $v$ is the reaction rate, $V_{max}$ is the maximal reaction rate, $K_m$ is the Michaelis-Menten constant, [S] is the substrate concentration, $K_i$ is the dissociation constant of the inhibitory enzyme-substrate- (SES) complex. For $IC_{50}$, and $EC_{50}$ determination, data were fitted to the corresponding dose-response equation. Data were analyzed with

GraphPad Prism 7.0 (GraphPad software, San Diego, CA).

### ATPase assay

For the DDX ATPase assay, Malachite Green Phosphate Assay Kit (MAK307, Sigma) was used according to manufacturer instructions. PolyA$^-$-RNA was obtained from *X. tropicalis* embryos as unbound material after mRNA purification. Briefly, embryo lysate was treated with proteinase K and total DNA/RNA was purified by Phenol/Chloroform extraction. RNA was separated from DNA by LiCl precipitation, passed through an oligo-dT column and the flow through was collected to yield PolyA$^-$-RNA. Reactions were carried out for 30 minutes at room temperature with 500 nM DDX proteins and 10 ng/µl PolyA$^-$-RNA in ATPase Buffer (30 mM HEPES-KOH pH 7.7, 50 mM KCl, 7 mM $MgCl_2$, 0.1 % BSA, 1 mM DTT, 0.1 mM EDTA, 5 % glycerol). Specific activity was expressed as pmol of phosphate released from ATP per minute.

### RNA filter binding assay

PolyA$^-$-RNA were radiolabeled with $^{32}$P-γATP by T4 PNK (NEB) following the manufacturer's instructions and quantified by UV spectrophotometry. 500 nM DDX or GFP control protein was incubated with increasing concentrations of PolyA$^-$-RNA for 15 minutes at 30 °C in ATPase Buffer. Reactions were spotted onto a bio-dot SF (Biorad) assembled from bottom to top as follows: 4 x 3MM paper, Nylon$^+$ membrane, and Nitrocellulose membrane all equilibrated in ATPase Buffer without BSA for 10 minutes. Wells were washed 5x with 150 µl ATPase Buffer without BSA, membranes were dried, and radioactivity was detected by phosphorimaging.

### ADP filter binding assay

500 nM DDX or GFP control protein was incubated without or with 1 µCi $^{32}$P-αADP (Hartmann Analytic SCP-227, 6000Ci/mmol) for 15 minutes at 30 °C in ATPase Buffer. Reactions were spotted onto a bio-dot SF (Biorad) and processed as described above for RNA filter binding assay.

### Thermal stability assay

The thermal stability of proteins was determined by nano differential scanning fluorimetry (nanoDSF), using a Prometheus NT.48 nanoDSF (NanoTemper technologies). 10 µM of each protein in Storage Buffer was loaded in standard capillaries (NanoTemper technologies).

Temperature was varied from 20 to 90 ºC, at 1 ºC/min rate. The fluorescence ratio 330nm/350nm was recorded, and the inflection point taken as melting temperature (Tm).

### phospho-HSP90 analysis

*Xenopus* embryos were injected at stage 2 - 3 with either 20 ng control or *ddx1* Morpholino (*X. tropicalis*) for loss-of-function studies, or with 500 pg *ck2*α and *ck2*β mRNA and/or 250 pg *ddx1* mRNA for gain-of-function studies (*X. laevis*). Embryos were collected at stage 18 and lysed in 4 µl (*X. tropicalis*) or 10 µl (*X. laevis*) Embryo Lysis Buffer (20 mM Tris pH 7.4, 150 mM NaCl, 2 % NP-40). Lysates were cleared by Freon extraction (CFC-113, Honeywell 34874), followed by centrifugation (14,000 rpm, 10 min at 4 °C), and analyzed by western blot. HeLa cells were seeded in 24-well plates (50.000 per well) and transfected with siRNA at a final concentration of 50 nM using Dharmafect1 (Dharmacon). Three days post transfection, cells were harvested in 100 µl HEK-Lysis Buffer, cleared by centrifugation, and analyzed by western blot.

### Sample preparation for MS

CoIP proteins were subjected to SDS-PAGE and stained with Quick Coomassie® (Serva). The gel lanes were cut into three slices containing the entire area of the resolved proteins and subjected to in-gel digestion for mass spectrometric analysis as previously described[38].Gel slices were destained followed by subsequent reduction, alkylation and overnight trypsin/LysC digestion. The supernatant containing the peptides was vacuum-dried and subsequently stored in 0.1% formic acid at -20 °C until LC-MS/MS analysis.

### LC-MS/MS analysis

Peptides were resolved using the Easy NanoLC1200 fitted with a trapping (Acclaim Pepmap C18, 5 µm, 100 A, 100 µm x 2 cm) and an analytical column (Acclaim PepMap RSLC C18, 2 µm, 100 A, 75 µm x 50 cm). The outline of the analytical column was coupled directly to a Fusion Orbitrap (Thermo Fisher Scientific) mass spectrometer. Solvent A was 0.1 % formic acid (vol/vol) and solvent B 80 % acetonitrile (vol/vol), 0.1 %formic acid (vol/vol). The peptides were loaded on the trap column with a constant flow of solvent A at a maximum pressure of 800 bar. Peptides were eluted from the analytical column at a constant flow rate of 300 nl/min and a temperature of 55 °C. During the elution the percentage of solvent B was increased in a linear gradient from 3 % to 8 % in 4 minutes, then from 8 % to 10 % in 2 minutes, then from 10 % to 32 % for 68 minutes and then from 32 % to 50 % for another 12 minutes. At the end of the gradient solvent B was kept at 100 % for 7 minutes followed by re-equilibration of the analytical column for 10 minutes at 97 % solvent A. The peptides were introduced to the mass spectrometer via a Pico-Tip Emitter 360 µm OD x 20 µm ID; 10 µm tip (New Objective) and a nano-source spray voltage of 2 kV. The ion transfer tube temperature was set to 275 °C. Full scan MS spectra 10 were acquired within the range (m/z) of 375 - 1500 in the Orbitrap detector with a resolution of 120,000. The maximum injection time was set to 50 ms and automatic gain control target (AGC) to $1x10^6$ ions. The most abundant ions within a 3 sec cycle time window were selected for fragmentation. Ions with unassigned charges and charges of 1 or $> 5$ were excluded. Dynamic exclusion was set to 40 sec with a mass tolerance of $\pm$10ppm. For the $MS^2$ scans the quadrupole was used with an isolation window of 1.6 m/z. For peptide fragmentation higher-energy collisional dissociation (HCD) was used at 33 %. $MS^2$ scans were acquired in the linear ion trap that was operated in the rapid ion scan rate with an AGC target of $1x10^4$ ions or a maximum injection of 50 ms. $MS^2$ scans were acquired as centroid data type. The mass spectrometry proteomics data have been deposited to the ProteomeXchange Consortium via the PRIDE partner repository with the dataset identifier PXD019405.

### Analysis of MS data

Raw files were processed using Maxquant software[39].(version 1.5.1.2). The search was performed against the *Xenopus laevis* uniprot canonical database (02/2016) containing both reviewed and TrEMBL entries. Enzyme digestion in Maxquant settings was set to Trypsin allowing for maximum of up to 2 missed-cleavages. Protein N-term acetylation, methionine oxidation were set as variable modifications and carbamidomethylation of cysteine as a fixed modification. Minimum unique peptides option was set to 1. Both intensity-based absolute quantification and label free quantification values were calculated. Peptide and protein hits were filtered at a false discovery rate (FDR) of 1 % with a minimal peptide length of 7 amino acids. The reversed sequences of the target database were used as a decoy database. Second peptide search for the identification of the chimeric $MS^2$ spectra was enabled. All other Maxquant options were left to their default settings. Subsequent analysis of the output tables was performed in Microsoft Excel and in R statistical software environment (version 3.4.3).

### Equilibrium fluorescence measurements

The fluorescence spectra of 250 µM mant-ADP (Jena Bioscience, cat. #NU-201S) were measured in the absence and presence of 2.5 µM CK2α2 using a Spectramax M5 spectrophotometer (Molecular Devices). Samples were prepared in a black 384-well flat-bottom plate (Greiner). Spectra were measured by exciting the samples at 290 nm and collecting the emission intensities between 390 and 500 nm. All steady-state fluorescence measurements were recorded at room temperature in Reaction Buffer (20 mM HEPES pH 8.0, 150 mM NaCl, 10 mM $MgCl_2$).

### Stopped-flow kinetic measurements

Kinetic measurements for monitoring the release of mant-ADP from CK2α2 were performed using a stopped-flow device (SX.18 MV, Applied Photophysics). 1 µM CK2α2 was preincubated with 20 µM mant-ADP, and then rapidly mixed in a 1:1 ratio with a solution containing 5 mM non-labeled ADP and either DDX27$^{215\text{-}651}$ or Ovalbumin (Sigma, #A5503 ) at indicated concentrations. All measurements were done at 10 °C in Reaction Buffer (20 mM HEPES pH 8.0, 150 mM NaCl, 10 mM $MgCl_2$ ). The excitation wavelength was 290 nm, and the emission was measured using a 420 nm cutoff filter. 10,000 datapoints were collected within 0.5 seconds. Each measurement is an average of at least 6 traces. To obtain rate constants, data were fitted to an exponential curve, either corresponding to a single or double decay, using the GraphPad Prism 7 software. The fluorescence decay for CK2α2 was biphasic. Analysis was limited to the fast phase. The slow phase was excluded, because it was unaffected by DDX27, and it might reflect off-pathway isomerization, or interactions with mant-ADP isoforms or impurities.

### Statistical analysis

All experiments, apart from kinetic experiments, were carried out in triplicates and repeated at least twice, except were indicated. GraphPad Prism 7.0 (GraphPad software, San Diego, CA) was used for all statistical analysis. One-way or two-way analysis of variance with Tukey's or Dunnett's multiple comparison test was used to determine statistical significance. Data are represented as mean ± standard deviation (SD) and data indicated as statistically significant with the following P-values: * $P \leqq 0.05$, ** $P \leqq 0.01$, *** $P \leqq 0.001$, **** $P \leqq 0.0001$, ns, no significant difference.

### Mathematical modeling, parameter inference and model selection

Enzyme-kinetic models for CK1ε and CK2α2 were developed based on ordinary differential equations and the quasi-steady-state assumption. Model parameters in the absence of DDX proteins were estimated by fitting models simultaneously to a large array of experimental data by the maximum-likelihood approach. Model selection addressing the mechanism of action of DDX proteins was performed by implementing best-fit parameters obtained in the absence of DDX proteins and estimating DDX effect sizes on different elementary steps of the enzyme-kinetic mechanisms, using maximum likelihood with experimental data measured in the presence of various DDX proteins. Details for modeling and model selection are provided in Supplementary Text. The Levenberg Marquardt algorithm was employed for optimization, with $> 104$ initial conditions for model parameters to avoid trapping in local minima. Custom computer programs for these tasks were coded in Python and will be made available upon request.

### Supplementary Text

#### Considerations for modeling enzymatic activity of kinases

Protein kinases catalyze the transfer of a phosphoryl group from the co-substrate ATP to an appropriate amino acid residue. Typically, this occurs by a ternary-complex mechanism (as opposed to a ping-pong mechanism), where both substrates are bound simultaneously. The formation of a ternary kinase-ATP-peptide substrate complex, may require a specific order of substrate binding (ordered binding), or may occur randomly. Ordered binding can be considered as a special case of a random mechanism. In the ternary complex, phosphoryl transfer yields the ternary kinase-ADP-phospho-peptide product complex, from which the reaction products are released. Importantly, phosphoryl transfer is a rapid process whose equilibrium configuration favors the ternary product complex, i.e., the equilibrium constant is much larger than 1[19]. As a consequence, dissociation of substrates from the ternary complex can be neglected.

Enzymatic two-substrate reactions, including those catalyzed by protein kinases, frequently display substrate inhibition—with increasing substrate concentration, the reaction velocity reaches a maximal value at some finite, critical substrate concentration, followed by a monotonic decline. This is typically due to at least one of the four reactants binding to the wrong enzyme species, to form a dead-end complex[19]. For example, if formation of the ternary substrate complex is ordered, with ATP binding before peptide, the binary kinase-peptide complex constitutes a dead-end. Another candidate for a substrate inhibition is a ternary kinase-ADP-peptide complex. Here, dissociation of the phospho-peptide from the ternary product complex results in a kinase-ADP complex, which can associate with a peptide. Then substrate inhibition will occur if ADP can only be released when no peptide is bound to the kinase, i.e., the ternary complex constitutes a dead-end for ADP release.

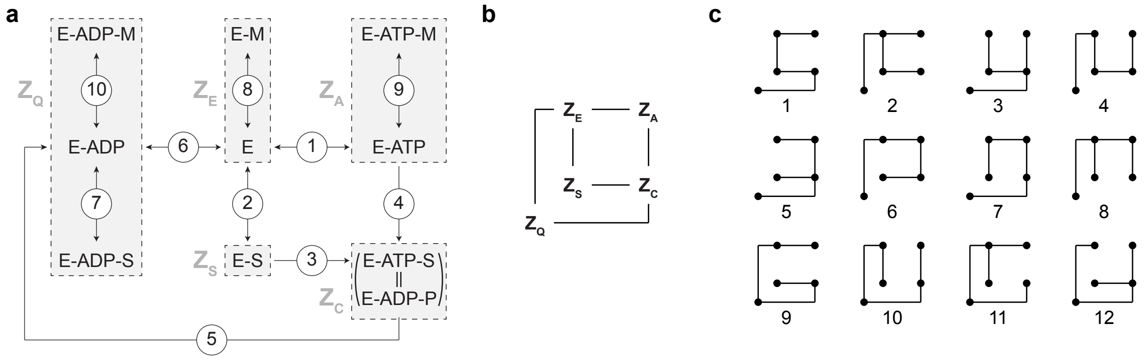


**Supplementary Text Figure** 1: **Derivation of reaction velocities.** (**a**) Reaction scheme as in Fig. 4a, with elementary steps numbered. The gray boxes indicate the effective reaction scheme used to derive the analytical expression for the reaction velocity. (**b**) Network representation of the effective reaction scheme. (**c**) The twelve King-Altman patterns of the effective scheme.

### Mathematical model

We develop a generic model of kinase kinetics that includes the possibility for both of the above-described mechanisms of substrate inhibition (binary and ternary dead-end complexes; Supplementary Text Fig. 1a). As explained above, substrate dissociation from the ternary complex is neglected as a consequence of rapid phosphoryl transfer. Moreover, we include binding of a non-phosphorylatable mutant peptide as a competitive inhibitor.

The numbering of the elementary reactions in Supplementary Text Fig. 1a defines the nomenclature for model parameters. Specifically, $k_n$ with positive subscript denotes the rate constant of reaction $n$ in direction of the reaction flux, while $k_{-n}$ denotes the rate constant of reaction $n$ in the reverse direction. The affinity of reaction $n$ is denoted with an upper case $K_n$. For example, the association rate of ATP binding to free kinase (reaction 1 in Supplementary Text Fig. 1a) is given by $k_1$, the corresponding dissociation rate by $k_{-1}$, and the affinity by $K_1 = k_1/k_{-1}$. For ADP binding a free kinase (reaction 6 in Supplementary Text Fig. 1a), the association rate is $k_{-6}$, the dissociation rate is $k_6$, and the affinity is $K_6 = k_{-6}/k_6$.

We use the King-Altman method to find the reaction velocity[40]. Let $A$, $S$, $Q$, and $M$ be the concentrations of ATP, peptide substrate, ADP, and mutant peptide, respectively. Since dead-end complexes, which are formed by reactions 7, 8, 9, and 10 (Supplementary Text Fig. 1a), obey the principle of detailed balance under steady-state kinetics, the effective reaction scheme, indicated by the gray boxes, suffices to derive an expression for the reaction velocity. This reaction scheme is translated into a network comprising

**Supplementary Table** 3: Contribution of each King-Altman pattern (Supplementary Text Fig. 1c) to each state of the effective reaction scheme (Supplementary Text Fig. 1b).

| King-Altman pattern | $Z_{\mathrm{E}}$ | $Z_{\mathrm{A}}$ | $Z_{\mathrm{S}}$ | $Z_{\mathrm{C}}$ | $Z_{\mathrm{Q}}$ |
|---|---|---|---|---|---|
| 1 | 0 | 0 | 0 | 0 | $k_{-1}k_2k_3k_5AS$ |
| 2 | 0 | 0 | 0 | $k_{-1}k_2k_3k_6AS$ | 0 |
| 3 | 0 | 0 | 0 | 0 | $k_2k_3k_4k_5AS^2$ |
| 4 | 0 | 0 | 0 | $k_2k_3k_4k_6AS^2$ | 0 |
| 5 | 0 | 0 | 0 | 0 | $k_1k_3k_4k_5A^2S$ |
| 6 | 0 | 0 | 0 | $k_1k_3k_4k_6A^2S$ | 0 |
| 7 | 0 | 0 | 0 | 0 | $k_1k_{-2}k_4k_5AS$ |
| 8 | 0 | 0 | 0 | $k_1k_{-2}k_4k_6AS$ | 0 |
| 9 | $k_{-1}k_3k_5k_6A$ | $k_1k_3k_5k_6A^2$ | 0 | 0 | $k_{-1}k_3k_5k_{-6}AQ$ |
| 10 | $k_{-2}k_4k_5k_6S$ | 0 | $k_2k_4k_5k_6S^2$ | 0 | $k_{-2}k_4k_5k_{-6}SQ$ |
| 11 | $k_{-1}k_{-2}k_5k_6$ | $k_1k_{-2}k_5k_6A$ | $k_{-1}k_2k_5k_6S$ | 0 | $k_{-1}k_{-2}k_5k_{-6}Q$ |
| 12 | $k_3k_4k_5k_6AS$ | 0 | 0 | 0 | $k_3k_4k_5k_{-6}ASQ$ |

five vertices and six edges (Supplementary Text Fig. 1c), for which twelve King-Altman patterns exist (Supplementary Text Fig. 1c), representing complete sub-networks with one edge less than the number of vertices and not containing any loops. Each King-Altman pattern delivers terms to the reaction velocity, as summarized in Supplementary Table 3. The reaction velocity $v$, normalized by the total kinase concentration $E_{\mathrm{tot}}$, is then given by

$$\frac{v}{E_{\mathrm{tot}}} = \frac{k_5 Z_{\mathrm{C}}}{Y}, \tag{2a}$$

$$Y \equiv (1 + K_8 M)Z_{\mathrm{E}} + (1 + K_9 M)Z_{\mathrm{A}} + Z_{\mathrm{S}} + Z_{\mathrm{C}} + (1 + K_7 S + K_{10} M)Z_{\mathrm{Q}}. \tag{2b}$$

Here, the $Z$'s represent the sum over all entries in the corresponding column of Supplementary Table 3, and the denominator of the reaction velocity, $Y$, is given by the sum over all entries of the table, with the prefactors accounting for the dead-end complexes. We introduce effective affinities for reactions 3 and 4,

$$K_{32} \equiv \frac{k_3}{k_{-2}}, \tag{3a}$$

$$K_{41} \equiv \frac{k_4}{k_{-1}}, \tag{3b}$$

as well as the following quantities,

$$X_{\mathrm{A}} \equiv 1 + K_{32}A, \tag{3c}$$

$$X_{\mathrm{S}} \equiv 1 + K_{41}S, \tag{3d}$$

$$X_{\mathrm{Q}} \equiv 1 + K_7 S + K_{10}M, \tag{3e}$$

with which we find for the denominator

$$\begin{aligned} Y = {} & (k_5 X_{\mathrm{Q}} + k_6)\,(k_3 K_2 X_{\mathrm{S}} + k_4 K_1 X_{\mathrm{A}})\,A\,S \\ & + k_5 k_6 \big[(1 + X_{\mathrm{Q}} K_6 Q + K_8 M) X_{\mathrm{A}} X_{\mathrm{S}} + (1 + K_9 M) X_{\mathrm{A}} K_1 A + X_{\mathrm{S}} K_2 S\big], \end{aligned} \tag{3f}$$

and for the reaction velocity,

$$\frac{v}{E_{\mathrm{tot}}} = \frac{k_5 k_6 (k_3 K_2 X_{\mathrm{S}} + k_4 K_1 X_{\mathrm{A}})\,A\,S}{Y}, \tag{4}$$

as a function of the kinetic parameters and concentrations of peptide substrate, ATP, ADP and non-phosphorylatable mutant peptide.

### Parameter estimation

Using the maximum likelihood approach, we fitted Eq. (4) to measured reaction velocities, obtained for various concentrations of peptide, ATP, ADP and mutant peptide (cf. Extended Data Fig. 5c for CK1ε, Extended Data Fig. 6c for CK2α2). We assumed that the measurement error follow a Gaussian distribution with a uniform coefficient of variation among all data. The resulting uncertainties in parameter estimates were computed with the profile likelihood method[41].and expressed as 95% confidence intervals (cf. Extended Data Fig. 5b for CK1ε, Extended Data Fig. 6b for CK2α2). For CK1ε, we found no significant contribution of reaction 3 (Extended Data Fig. 5a,b), indicating ordered substrate binding with ATP preceding peptide.

### Modeling the effect of DDX proteins on kinase activity

**Mathematical model**. To account for the effect of DDX protein on kinase activity, we let DDX proteins impact one or more rate constants of the enzyme-kinetic model. We then use model selection against reaction velocity data measured in the presence of various DDX proteins to find parsimonious models for DDX action on CK1εand CK2α2. Specifically, DDX proteins may act on association and dissociation rates of substrates and products. We allow for independent DDX effects on the peptide and nucleotide binding sites. For a given site, all chosen rates (dissociation and/or association) are altered by the same factor for a specific DDX protein. For example, consider nucleotide association, for which there are the association rates of ATP binding, $k_1$ and $k_3$, and the association rate of ADP binding, $k_{-6}$. We define the DDX effect size on nucleotide association, $\delta_{\mathrm{nuc,on}}$, via

$$\begin{aligned} k_1' &= \delta_{\mathrm{nuc,on}}\, k_1, \\ k_3' &= \delta_{\mathrm{nuc,on}}\, k_3, \\ k_{-6}' &= \delta_{\mathrm{nuc,on}}\, k_{-6}. \end{aligned} \tag{5}$$

with the $k'$s now appearing in Eqs. (3) and (4). Corresponding definitions hold for the DDX effect on the dissociation rates of nucleotide binding, $\delta_{\mathrm{nuc,off}}$, as well as the association and dissociation rates of peptide binding, $\delta_{\mathrm{pep,on}}$ and $\delta_{\mathrm{pep,off}}$.

**Determining the effect of DDX proteins**. For each binding site of the kinase we consider four scenarios: DDX impacts (i) only association, (ii) only dissociation, (iii) both association and dissociation with the same magnitude, and (iv) a DDX effect is absent. Scenarios (i) and (ii) change the affinity of the respective site whereas scenario (iii) models a purely kinetic effect. Thus, there are sixteen possible combinations of DDX effects. The resulting models were tested against a comprehensive data set, comprising fold-changes in reaction velocity in the presence of DDX proteins, measured under rate-limiting conditions for ATP binding, peptide binding, and product release for each kinase and DDX protein. We fitted the models with the maximum likelihood method, using the best-fit parameters for the enzyme-kinetic models without DDX protein and estimating the DDX effect sizes (the $\delta$ parameters introduced above). We assume homoscedasticity for measurement uncertainties and compared the models using the Akaike Information Criterion[20] (see Fig. 4g).

**Dose dependence**. The dose dependence of the DDX effect size on reaction group $j$ is described by a characteristic dose $D^*$ and a maximal effect size $\bar{\delta}_j$,

$$\delta_j(D) = \frac{D_j^* + \bar{\delta}_j D}{D_j^* + D}. \tag{6}$$

Note that $\delta_j(0) = 1$ and $\delta_j = \bar{\delta}_j$ as $D \to \infty$.

### Supplementary References


33. Bogomolovas, J., Simon, B., Sattler, M. & Stier, G. Screening of fusion partners for high yield expression and purification of bioactive viscotoxins. Protein Expr Purif 64, 16-23 (2009).

34. Dominguez, I. et al. Protein kinase CK2 is required for dorsal axis formation in Xenopus embryos. Dev Biol 274, 110-24 (2004).

35. Acebron, S.P., Karaulanov, E., Berger, B.S., Huang, Y.L. & Niehrs, C. Mitotic wnt signaling promotes protein stabilization and regulates cell size. Mol Cell 54, 663-74 (2014).

36. Tomishima, M.J., Hadjantonakis, A.K., Gong, S. & Studer, L. Production of green fluorescent protein transgenic embryonic stem cells using the GENSAT bacterial artificial chromosome library. Stem Cells 25, 39-45 (2007).

37. Copeland, R. Enzymes: a practical introduction to structure, mechanism, and data analysis. (2000).

38. Shevchenko, A., Tomas, H., Havlis, J., Olsen, J.V. & Mann, M. In-gel digestion for mass spectrometric characterization of proteins and proteomes. Nat Protoc 1, 2856-60 (2006).

39. Tyanova, S., Temu, T. & Cox, J. The MaxQuant computational platform for mass spectrometry-based shotgun proteomics. Nat Protoc 11, 2301-2319 (2016).

40. King, E.L. & Altman, C. A Schematic Method of Deriving the Rate Laws for Enzyme-Catalyzed Reactions. The Journal of Physical Chemistry (1956).

41. Venzon, D.J. & Moolgavkar, S.H. A Method for Computing Profile-Likelihood-Based Confidence Intervals. Applied statistics, 37, 87-94. (1988).
